# Unravelling the Transcriptomic Symphony of Sarcopenia: Key Pathways and Hub Genes Altered by Muscle Ageing and Caloric Restriction Revealed by RNA Sequencing

**DOI:** 10.1101/2024.03.09.584213

**Authors:** Gulam Altab, Brian J. Merry, Charles W. Beckett, Priyanka Raina, Inês Lopes, Katarzyna Goljanek-Whysall, João Pedro de Magalhães

## Abstract

Sarcopenia is a disease involving extensive loss of muscle mass and strength with age and is a major cause of disability and accidents in the elderly. Mechanisms purported to be involved in muscle ageing and sarcopenia are numerous but poorly understood, necessitating deeper study. Hence, we employed high-throughput RNA sequencing to explicate the global changes in protein-coding gene expression occurring in skeletal muscle with age. Caloric restriction (CR) is a proven prophylactic intervention against sarcopenia. Therefore, total RNA was isolated from the muscle tissue of both rats fed ad libitum and CR rats. Collected data were subjected to Gene Ontology, pathway, co-expression, and interaction network analyses. This revealed the functional pathways most activated by both ageing and CR, as well as the key “hub” proteins involved in their activation.

RNA-seq revealed 442 protein-coding genes to be upregulated and 377 to be downregulated in aged muscle, compared to young muscle. Upregulated genes were commonly involved in protein folding and the immune response; meanwhile, downregulated genes were often related to developmental biology. CR was found to suppress 69.7% and rescue 57.8% of the genes found to be upregulated and downregulated in aged muscle, respectively. In addition, CR uniquely upregulated 291 and downregulated 304 protein-coding genes. Hub genes implicated in both ageing and CR included *Gc*, *Plg*, *Irf7*, *Ifit3*, *Usp18*, *Rsad2*, *Blm* and *RT1-A2*, whilst those exclusively implicated in CR responses included *Alb*, *Apoa1*, *Ambp*, *F2*, *Apoh*, *Orm1*, *Mx1*, *Oasl2* and *Rtp4*. Hub genes involved in ageing but unaffected by CR included *Fgg*, *Fga*, *Fgb* and *Serpinc1*. In conclusion, this comprehensive RNA sequencing study highlighted gene expression patterns, hub genes and signalling pathways most affected by ageing in skeletal muscle. This data may provide the initial evidence for several targets for therapeutic interventions against sarcopenia.

## Introduction

Sarcopenia is an age-related disease, defined as a “loss of walking speed or grip strength associated with low muscle mass” and is said to affect around 10% of those over 65 and half of those over 80 ^1^. Since its inclusion in the World Health Organisation’s revision of the International Statistical Classification of Diseases and Related Health Problems in 2016 (ICD-11), there has been a surge of research interest in the disease ^2,3^. However, the causes of muscle ageing are complex and enigmatic. Implicated mechanisms include satellite cell exhaustion ^4,5^, reduced proteostasis capacity ^6^, increased ROS production ^7^, increased inflammation ^8^, and mitochondrial dysfunction ^9^. Understanding both the mechanisms involved in the ageing process, as well as how it may be ameliorated, is crucial to improving the quality of life of elderly patients affected by sarcopenia.

Given its high degree of network complexity, -omics approaches are likely best placed to explain the underlying causal structure of muscle ageing ^10^. Previous studies have assessed the transcriptomic signature of muscle ageing in mice ^11–13^ but only one has done so in rats ^14^. Caloric restriction (CR) has repeatedly been demonstrated to be an effective prophylactic against age-related muscle mass loss in laboratory rodents ^15–17^ and non-human primates ^18–20^. No studies to our knowledge, however, have assessed the transcriptomic correlates of CR-mediated delay of muscle ageing in rats. Using rat tissue, we aimed to replicate previous findings regarding the key changes in protein-coding gene expression observed in muscle ageing. In addition, we endeavoured to assess to what extent these changes could be reversed using CR. Furthermore, we employed Gene Ontology (GO) and Reactome pathway analyses to assess in which functional pathways identified genes were implicated. Finally, we used gene co-expression and protein-protein interaction (PPI) network analyses to identify the most important “hub” genes mediating the effects of both ageing and CR in rat muscle.

## Materials and methods

### Animals

Gastrocnemius muscle tissue samples were collected from rats used in a previous study conducted by ^21^, under Home Office Project License 40/2964. All animal experiments were conducted in compliance with the United Kingdom Animals (Scientific Procedures) Act of 1986. In brief, 21-28 day old, male, Brown Norway (Substrain BN/SsNOlaHSD) rats were acquired from Harlan Laboratories, UK. Animals were kept under barrier conditions on a 12-h light:12-h dark cycle (08:00– 20:00). Prior to 2 months of age, rats were group housed (n=4) and fed ad libitum (diet procured from Special Diet Systems Division Dietex International in Witham, UK). Following maturation, all animals were moved to solitary housing and randomly assigned to either ad libitum (AL) or caloric restriction (CR) diet groups. The CR rats were fed the same diet as the ad libitum rats, but their daily intake was restricted to 55% of the age-matched ad libitum intake. The CR rats were given access to food for a limited time each day between 10:30 and 11:00 hrs, while the ad libitum rats had unlimited access to the food for 24 hours per day^21^.

Animals were either terminated at 6 or 28 months of age (Table 1). No animals displayed any signs of clinical pathology prior to sacrifice. Following termination, gastrocnemius muscle tissue was removed, flash-frozen in liquid N_2_ and stored at −80 °C, prior to further analysis.

**Table 1.**
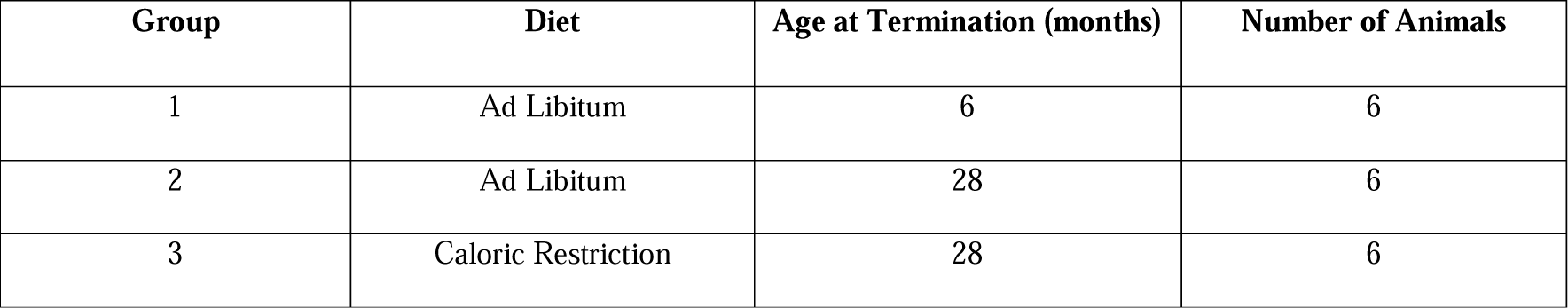
Group Identity, Diet Regimen and Age at Termination. Tissue samples were collected from male Brown Norway rats from one of 3 groups (n=6). Following animal arrival at 21-28 days of age, all animals were fed ad libitum (AL). After reaching full maturation/2-months old, animals were randomly allocated to either an AL (groups 1-2) or caloric restriction (CR) diet (group 3). Half the AL-fed animals were terminated at 6-months old and half at 28-months old. CR rats were terminated at 28-months old. Tissue samples were collected immediately post-sacrifice, snap frozen and stored at −80 °C until the day of analysis.

### RNA Extraction and RNA**□**Seq Analysis

Following removal from storage, gastrocnemius muscle tissues were immediately transferred onto ice to prevent thawing and degradation. Subsequently, each tissue was ground into a powder using a mortar and pestle. Liquid nitrogen was added periodically to ensure tissues remained in their frozen state. Total RNA was isolated using RNeasy Plus Universal Mini Kit (Qiagen, UK) according to manufacturer instructions. Extracted RNA quantity and quality were assessed on nanoDrop 2000 spectrophotometer and using the Agilent 2100 Bioanalyser, revealing all extracted RNA sampled to be of high quality (RINs>8). RNA integrity was further examined using agarose gel electrophoresis. Ribosomal RNA (rRNA) was removed from the total RNA using Illumina Ribo-Zero Plus rRNA depletion kit following the manufacturer’s instructions. Strand-specific RNA-seq library preparation was done using NEBNext® Ultra™ Directional RNA Library Prep Kit for Illumina, according to manufacturer protocol. Highlthroughput RNAlseq was performed at the Centre for Genomic Research at the University of Liverpool on the Illumina Hiseq 4000 platform (Paired-end, 2×150 bp sequencing, generating data from >280 M clusters per lane). The raw Fastq files were trimmed for the presence of Illumina adapter sequences using Cutadapt version 1.2.1. The reads were further trimmed using Sickle version 1.200 with a minimum window quality score of 20. Lastly, short reads <20 bp in length were removed.

The filtered reads were mapped to the rat reference genome (ftp.ensembl.org/pub/release-100/fasta/rattus_norvegicus/dna/) using STAR 2.3.0e. MultiQC showed the alignment score was above 80% for all the samples. To count the number of reads that mapped to each gene, featureCount was used to obtain the read count. Ensembl Rattus_norvegicus.Rnor_6.0.96.gtf were chosen as annotation references for the analyses.

### Differential Expression Analysis

Differentially expressed genes (DEGs) were identified by using the Bioconductor package “DESeq2” (version, 3.7) in R (version 4.1.0). Raw gene counts, detected by featureCount, were used as the input. DESeq2 is a well-known method, which offers statistical procedures for identifying differential expression by using negative binomial linear models for individual genes and uses the Wald test for significance analysis. The P-values from the Wald test were then adjusted using the Benjamini and Hochberg’s method for controlling the false discovery rate (p-adj). Differentially expressed genes were defined as those with a padj-value < =0.05 and fold change (FC)>=1.5. Biomart (https://www.ensembl.org/biomart) was used to annotate the biotype of each gene.

### Gene Ontology and Pathway Analysis

For the GO anslysis, we analysed distinct sets of differentially expressed genes separately. Initially, we compared gene expression profiles between young and old samples, delving into the functional implications of both upregulated and downregulated genes in the ageing process. Subsequently, we explored the differentially expressed genes between young and old samples under caloric restriction (CR), investigating their potential roles in ageing under CR conditions. Using Venn diagrams, we further classified these genes into specific groups: those exclusively upregulated or downregulated in either old or old CR muscle, genes suppressed by CR in old muscle, or genes uniquely differentially expressed in old CR muscle. For each subgroup, we conducted gene ontology analysis to elucidate their functional significance in the context of ageing and CR.

Both GO and Reactome pathway enrichment analyses were performed using ClusterProfiler (version 3.14.3), which enabled visualization of functional profiles ^22^. For this study, we performed GO biological process (GO-BP) analysis in the general level (GO level 3-8). Revigo (http://revigo.irb.hr) was used to remove redundant GO terms, clustering them based on semantic similarity. Input gene sets compared to the background by the hypergeometric test were used to obtain the significant GO terms and Reactome pathways (p<0.05) and the p-values corrected for false discovery rate (FDR)(padj) <=0.05 in multiple testing, q-values are also projected for FDR control. The top 20 GO and Reactome pathways were then visualised using ggplot2 (version 3.3.5) in R.

### Co-expression Network Construction

Using the weighted gene co-expression network analysis (WGCNA) package in R, an unsigned gene co-expression network was produced ^23^. Briefly, an adjacency matrix (Adj) was created from the input gene list by calculating Pearson correlations (Adj = |0.5 x (1 + Corr)| ß) between all pairs of genes among all the provided samples (n=6/ group). Adj was then converted to Topological Overlap Matrix (TOM) using the TOMsimilarity function in R. TOM represent the inter-connectedness between a pair of gene, which was then used to create a network tree through hierarchical clustering. Gene clusters with high topological overlap were designated as a module. Using the dynamicTreeCut R package, modules were detected from the dendrogram and were labelled with a standard colour scheme ^24^. The expression profile of genes in each module were represented by module eigengene (ME). The correlation between module eigengene and the gene expression profile was calculated to obtain module membership (MM), and the correlation between the gene profile and the trait (age, e.g., 6 months and 28 months old) was computed to attain Gene Significance (GS). Then, associations between module membership (MM) vs. gene significance (GS) were calculated. GS and MM values were then used to identify hub genes in modules that were strongly correlated with traits of interest, using stringent thresholds (GS values >0.80 and MM values >0.80). Finally, interesting modules were visualized by Cytoscape software and further analysed by protein-protein interaction (PPI) network analysis. Weighted Gene Correlation Network Analysis (WGCNA) was conducted using normalised counts of 835 differentially expressed genes from old rat muscle to identify hub genes associated with muscle ageing. A soft power of 22 was chosen, allowing the discovery of co-expression modules.

### Protein-Protein Interaction (PPI) Network Analysis

To construct PPI networks “The search tool for retrieval of interacting genes” (STRING) (https://string-db.org) was used ^25^. The STRING database allows the identification of known and predicted interaction between proteins, given genes of interest. A stringent threshold interaction score ofl >l0.7 was used to construct the PPI networks. Networks were downloaded and visualised in Cytoscape software (version 3.9.1). Using cytoHubba (version 0.1), a plugin of Cytoscape, hub proteins/genes were identified ^26^using 3 different methods, the maximal clique centrality (MCC) method, maximum neighbourhood component (MNC) method and Degree (Deg) method. This facilitated the the identification of the top ten hub genes associated with ageing and CR ^26^. Then, Venn diagrams were generated using Venny 2.1 (https://bioinfogp.cnb.csic.es/tools/venny/) to visualise which hub genes overlapped between the 3 methods ^27^. Lastly, the overlap between age-associated and CR-associated hub genes was assessed.

### RT-QPCR Validation of RNA-seq Data

To validate RNA-seq data results, reverse transcriptase quantitative PCR (RT-qPCR) analysis was used. Validation was restricted to a minimum of 3 samples per condition (n=>3) for whom muscle RNA was still available. Six RNAs that were greatly linked with ageing or dysregulated in old muscle, were chosen for validation. These included *Foxo1* (p-adjl=l0.0004), *Ogdh* (p-adjl=l0.0001), *Klf4* (pl=l0.005), *Pgc1a* (p-adjl=l0.001), *Cry1* (p-adj = 0.00001), and *Nr4a3* (p-adj = 0.000000002) mRNAs.

RNeasy Plus Universal Mini Kit (Qiagen, UK) was used according to the manufacturer’s protocol, to isolate the total RNA. Extracted RNA quantity and quality were assessed on the nanoDrop 2000 spectrophotometer and using the Agilent 2100 Bioanalyser. The miRCURY LNA RT Ki (Qiagen, UK) was used to synthesise cDNA and miRCURY LNA RT Kit (Qiagen, UK) was used to perform qPCR, according to manufacturer’s protocol in Roche ® LightCycler® 480. F-box protein 45 (Fbxo45) was identified as a relevant housekeeping gene using geNORM^28^ thus, used as an internal control for RNA template normalization, F: AAGTCAACGGCAGCTTCCCACA, R: CCAGGAACTCATACCCACGCTC (Origene, USA). Primers were obtained from previous literature or previously validated in our lab. *Foxo1*, F: AATTTGCTAAGAGCCGAGGA, R: AAGTCATCATTGCTGTGGGA ^29^. *Nr4a3*, F: AGACAAGAGACGTCGAAATCGAT, R: CTTCACCATCCCGACACTGA ^30^. *Ogdh*, F: GAGCTGAACAGGAGACAGGTAT, R: CGTCCTAATTGCTGCTGGTT. *Ppargc1a*, F: AGGTCCCCAGGCAGTAGAT, R: CGTGCTCATTGGCTTCATA ^31^. *klf4*, F: CAGCTGGCAAGCGCTACA, R: CCTTTCTCCTGATTATCCATTC ^32^. *Cry1*, TTCGCCGGCTCTTCCAA, R: ATTGGCATCAAGGTCCTCAAGA ^33^. The delta-delta Ct method (2–ΔΔCt method) was used to calculate the relative expression levels. The result mean ± SEM were plotted using Prism (version, 5.01) for Windows. One-way ANOVA (analysis of variance) and Tukey’s Multiple Comparison post-hoc test was conducted to compare the difference between the groups. The null hypothesis of no model effects was accepted at P<0.05.

## Results

### Age-related gene expression changes in the gastrocnemius muscle of AL-fed and CR rats

DESeq2 revealed 835 protein-coding genes to be differentially expressed in the gastrocnemius muscle of aged (28-month-old) rats fed AL when compared to young (6-month-old) rats fed AL. Of these, 442 genes were upregulated, whilst 377 were downregulated, in aged muscle, compared to young muscle. Genes with the greatest degree of up-regulation included Dclk1, Serpina12, *RT1-A2, Spg21, Igtp, Lgals1, B2m, Ctxn3, Bid* and *Nlrc5* (Table, 2A). Genes with the greatest degree of down-regulation included *Kcnc1, Nr4a2, Arntl, Nr4a3, Col1a1, Maf, Mpp3, AABR07044412.1, Auts2* and *Ddit4* (Table 2B).

**Table 2.**
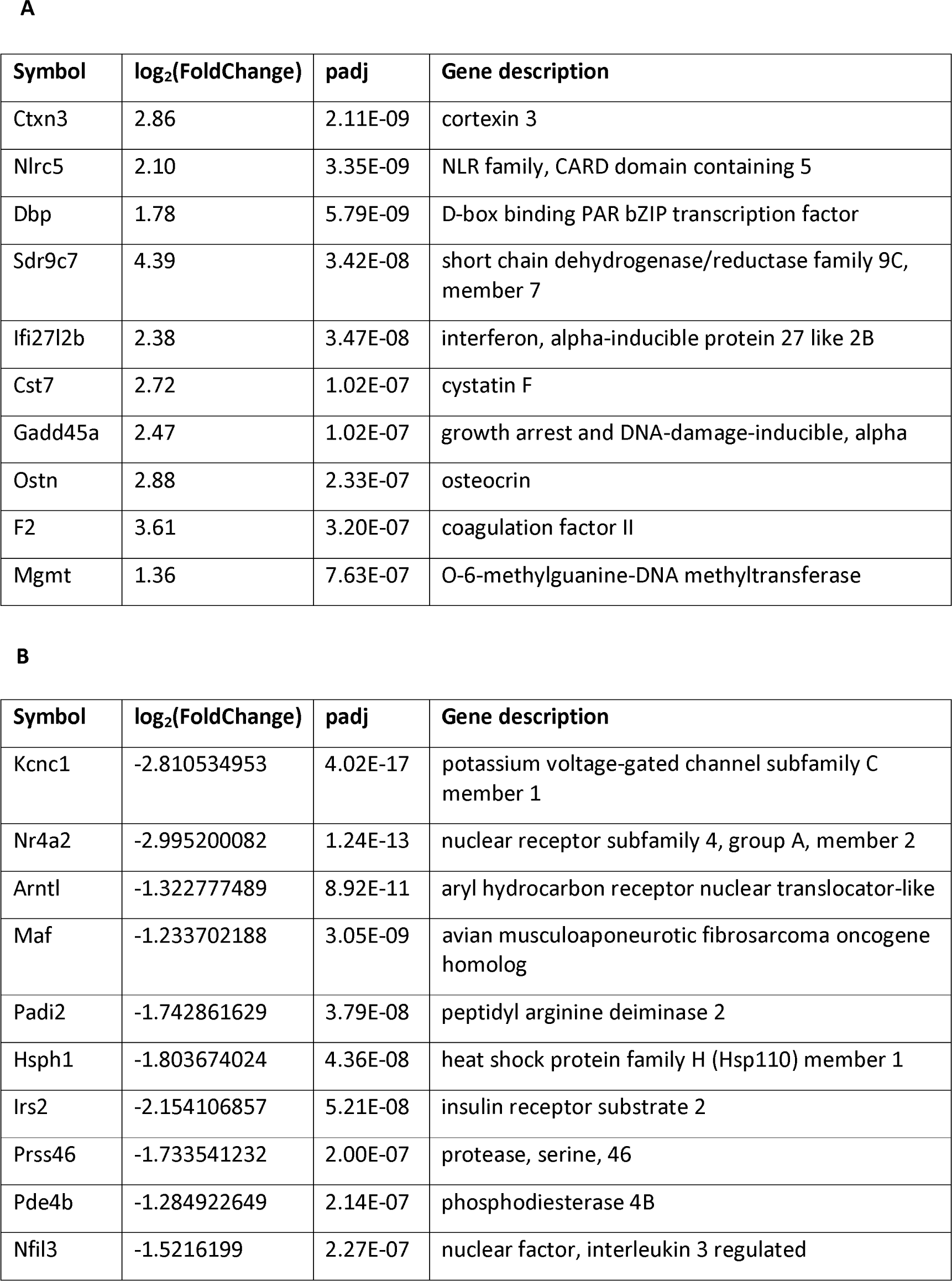
Genes with the largest magnitude of differential expression in the muscle of aged rats fed ad libitum (AL), when compared to the muscle of young rats fed AL. The tables list the gene most A. upregulated and B. downregulated by ageing. Differential expression was defined as a padj-value < =0.05 and fold change >=1.5. Approximate descriptions of gene function, revealed by gene ontology (GO) analysis, are listed in in the far-right column.

DESeq2 revealed 903 protein-coding genes to be differentially expressed in the gastrocnemius muscle of aged CR rats when compared to aged rats fed AL. Of these, 425 were upregulated, whilst 463 were downregulated, in aged muscle, compared to young muscle. Genes with the greatest degree of up-regulation included Dbp, Txndc16, *Cst7, Per3, Hook1, Cpt1a, Rusc2, Slc7a2, Tef* and *Adamts20* (Table, 3A). Genes with the greatest degree of down-regulation included *Arntl, Ubc, Slc45a3, Slc41a3, Npas2, Tead4, Asb2, Nfil3, AABR07044412.1* and *Dyrk2* (Table, 3B).

**Table 3.** Genes with the largest magnitude of differential expression in the muscle of aged calorically restricted rats when compared to the muscle of aged rats fed ad libitum. The tables list the gene most A. upregulated and B. downregulated by exposure to caloric restriction. Differential expression was defined as a padj-value < =0.05 and fold change >=1.5. Approximate descriptions of gene function, revealed by gene ontology (GO) analysis, are listed in in the far-right column.

**A**
| Symbol | log<sub>2</sub>(FoldChange) | padj | Gene description |
| --- | --- | --- | --- |
| B2m | 1.27 | 7.68E-10 | beta-2 microglobulin |
| Bid | 1.16 | 3.09E-09 | BH3 interacting domain death agonist |
| Dclk1 | 2.80 | 6.08E-20 | doublecortin-like kinase 1 |
| Igtp | 1.01 | 2.14E-11 | interferon gamma induced GTPase |
| Krt18 | 4.38 | 1.62E-08 | keratin 18 |
| Lgals1 | 0.78 | 5.60E-10 | galectin 1 |
| Mab21l3 | 3.84 | 4.35E-09 | mab-21 like 3 |
| Serpina12 | 4.02 | 4.83E-14 | serpin family A member 12 |
| Sh3kbp1 | 0.79 | 2.48E-08 | SH3-domain kinase binding protein 1 |
| Spg21 | 0.93 | 1.54E-12 | SPG21 abhydrolase domain containing, maspardin |

| Symbol | log<sub>2</sub>(FoldChange) | padj | Gene description |
| --- | --- | --- | --- |
| Col1a1 | -1.82 | 2.11E-09 | collagen type I alpha 1 chain |
| Ddit4 | -1.46 | 2.48E-08 | DNA-damage-inducible transcript 4 |
| Dusp8 | -1.21 | 5.36E-07 | dual specificity phosphatase 8 |
| Gpr157 | -1.59 | 4.84E-07 | G protein-coupled receptor 157 |
| Mpp3 | -1.91 | 4.50E-09 | membrane palmitoylated protein 3 |
| Mylk4 | -1.34 | 9.58E-08 | Myosin Light Chain Kinase Family Member 4 |
| Nr4a1 | -1.91 | 4.20E-08 | nuclear receptor subfamily 4, group A, member 1 |
| Nr4a3 | -2.86 | 1.59E-09 | nuclear receptor subfamily 4, group A, member 3 |
| RGD1565323 | -1.23 | 8.69E-07 | similar to OTTMUSP00000000621 |
| Slc25a25 | -1.92 | 7.89E-08 | solute carrier family 25 member 25 |

### Effect of CR on Age-related changes in gene expression in the rat gastrocnemius muscle

Comparison between the muscle of aged rats fed AL and aged CR rats, revealed CR to ameliorate age-related changes in the expression of 516 protein-coding genes (Figure 1). Of the genes found to be upregulated in aged compared to young rats fed AL, 308 (69.7%) were suppressed in aged CR rats, whilst 134 (30.3%) were unaffected. Of the genes found to be downregulated in aged compared to young rats fed AL, 218 (57.8%) were rescued in aged CR rats, whilst 159 (42.2%) were unaffected. In addition, 291 and 304 protein-coding genes were respectively upregulated and downregulated uniquely by CR.

**Figure 1.**
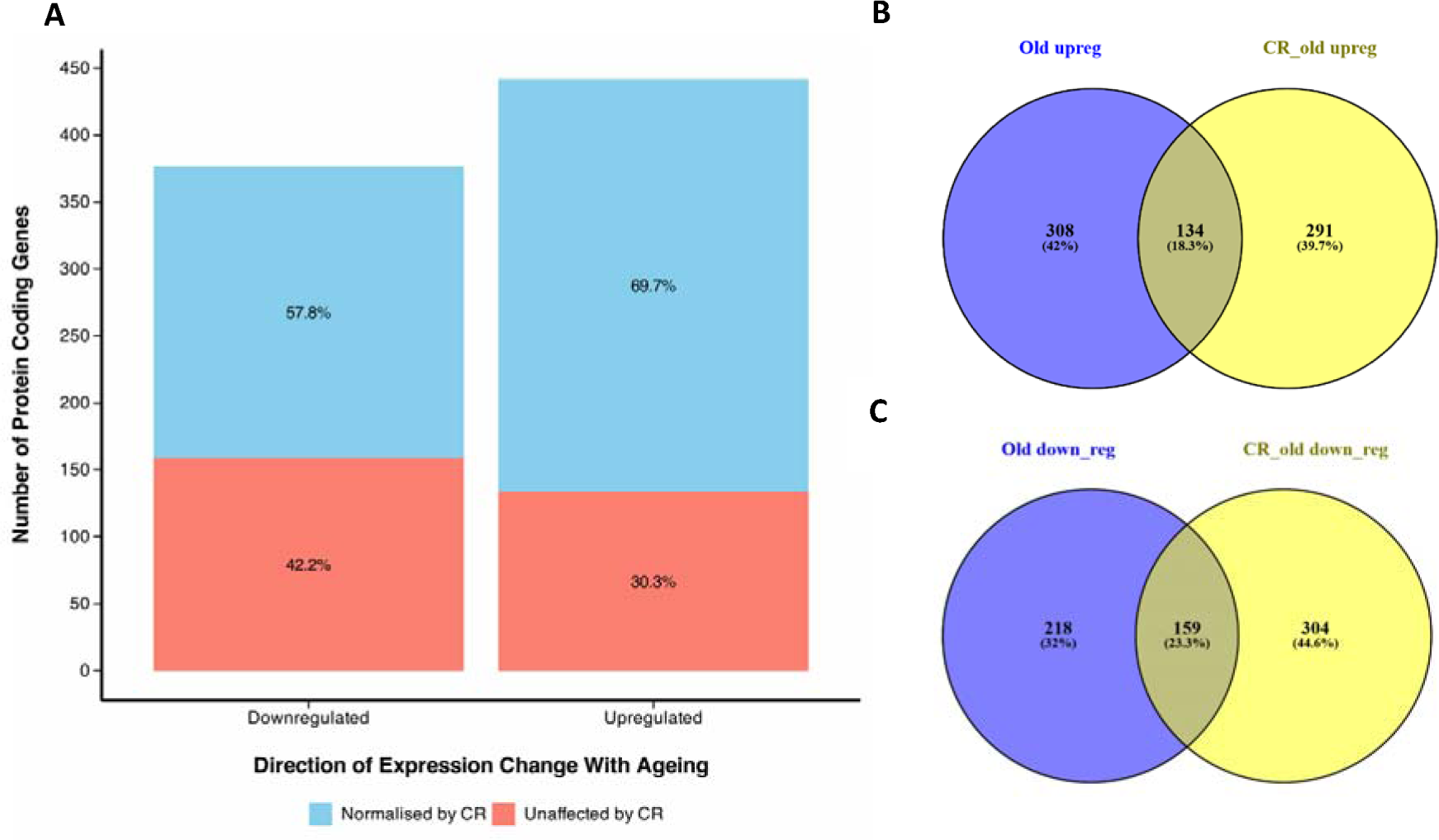
Impact of CR on gene expression. A. A bar chart showing the number of genes upregulated and downregulated in the muscle of aged rats fed ad libitum (AL), when compared to the muscle of young rats fed AL, and what percentage of these are normalised by caloric restriction (CR). Venn diagrams describe the number of genes B. upregulated and C. downregulated uniquely in old rats fed AL (purple), uniquely in old CR rats (yellow) and in both groups (centre), when compared to young rats fed AL. Differential expression was defined as FC 1.5-fold, and padj <=0.05. Venn diagrams were created using “ https://bioinfogp.cnb.csic.es/tools/venny/”.

### Functional and pathway analysis of protein-coding genes with differential expression in aged rats fed AL

ClusterProfiler gene ontology (GO)-analysis revealed 456 GO terms to be significantly associated with the genes upregulated in aged compared to young rats fed AL. Top upregulated terms based on number of genes included innate immune response, adaptive immune response and cell killing. Revigo summarised these upregulated genes to be primarily involved in lymphocyte-mediated immunity, steroid metabolism, cell killing, negative regulation of responses to external stimuli and coagulation (Figure 2A). Subsequently, upregulated genes were found to be significantly enriched for 48 Reactome pathways, including plasma lipoprotein remodelling, regulation of insulin-like growth factor (IGF) transport and uptake by binding proteins, post-translational protein phosphorylation, TNFR2 non-canonical, the NF-kB pathway and biological oxidations (Figure 2C).

**Figure 2.**
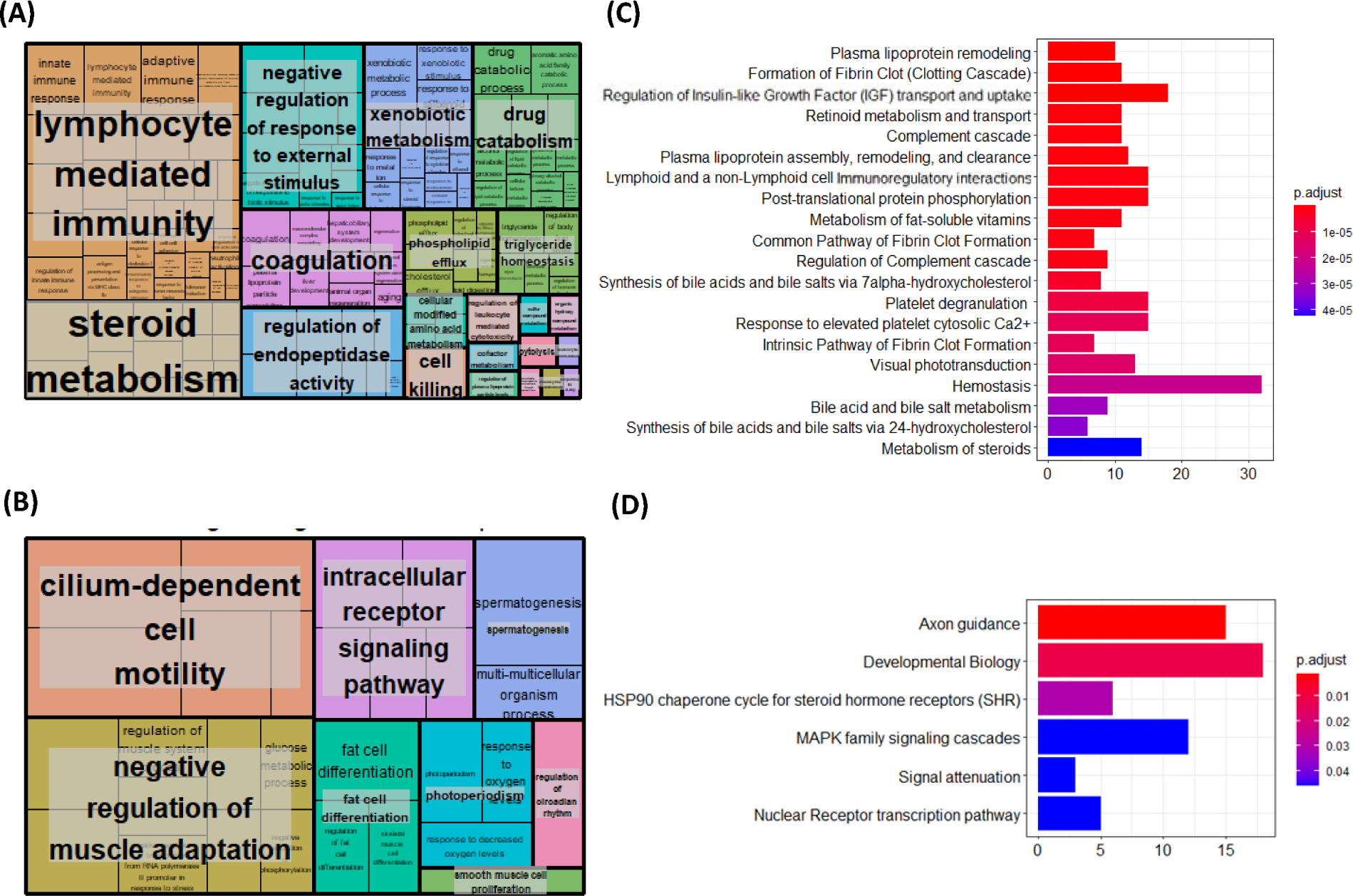
Gene Ontology (GO) and Reactome pathway analysis of genes differentially expressed downregulated in aged compared to young rat muscle. ClusterProfiler (version 3.14.3) package for R, was used to perform GO enrichment analysis. All the genes that were detect in skeletal muscle by RNA-seq analysis was used as background. A padj <0.05 was used as the cut off p-value. Revigo (http://revigo.irb.hr) then summarised GO terms for A. upregulated genes and B. downregulated genes. Subsequently, the ReactomePA (version 3.15) package was used for pathway analysis in R software (version 4.1), revealing Reactome pathways significantly (p-adj value=<0.05) associated with C. upregulated and D. downregulated genes.

ClusterProfiler revealed 40 GO terms to be significantly associated with the genes downregulated in aged compared to young rats fed AL. Top downregulated terms based on gene size included spermatogenesis, negative regulation of carbohydrate metabolism and negative regulation of skeletal muscle cell adaptation, fat cell differentiation, nucleotide catabolic process and skeletal muscle cell differentiation. Revigo summarised downregulated genes to primarily be involved in cilium-dependant motility, negative regulation of muscle adaptation, intracellular signalling and fat cell differentiation (Figure 2B). Subsequently, downregulated genes in old CR rats were found to be significantly enriched for 6 Reactome pathways. These pathways included axon guidance, developmental biology (including myogenesis), HSP90 chaperone cycle for steroid hormone receptors (SHR), MAPK family signalling cascades and nuclear receptor transcription pathways (Figure 2D).

### Functional and pathway analysis of protein-coding genes normalised by CR

ClusterProfiler revealed 265 GO terms to be significantly associated with genes suppressed in aged CR rats compared to age rats fed AL. Top terms based on gene size included innate immune response, response to molecule of bacterial origin and complement activation cascade. Revigo summarised genes suppressed by CR to primarily be involved in the innate immune response, triglyceride homeostasis and steroid metabolism (Figure 3A). Subsequently, genes suppressed by CR were found to be significantly enriched for 18 Reactome pathways, including the clotting cascade, plasma lipoprotein remodelling, plasma lipoprotein assembly, remodelling, and clearance, regulation of IGF transport and uptake by IGF binding proteins, retinoid metabolism and transport, complement cascade and post-translational protein phosphorylation (Figure 3C).

**Figure 3.**
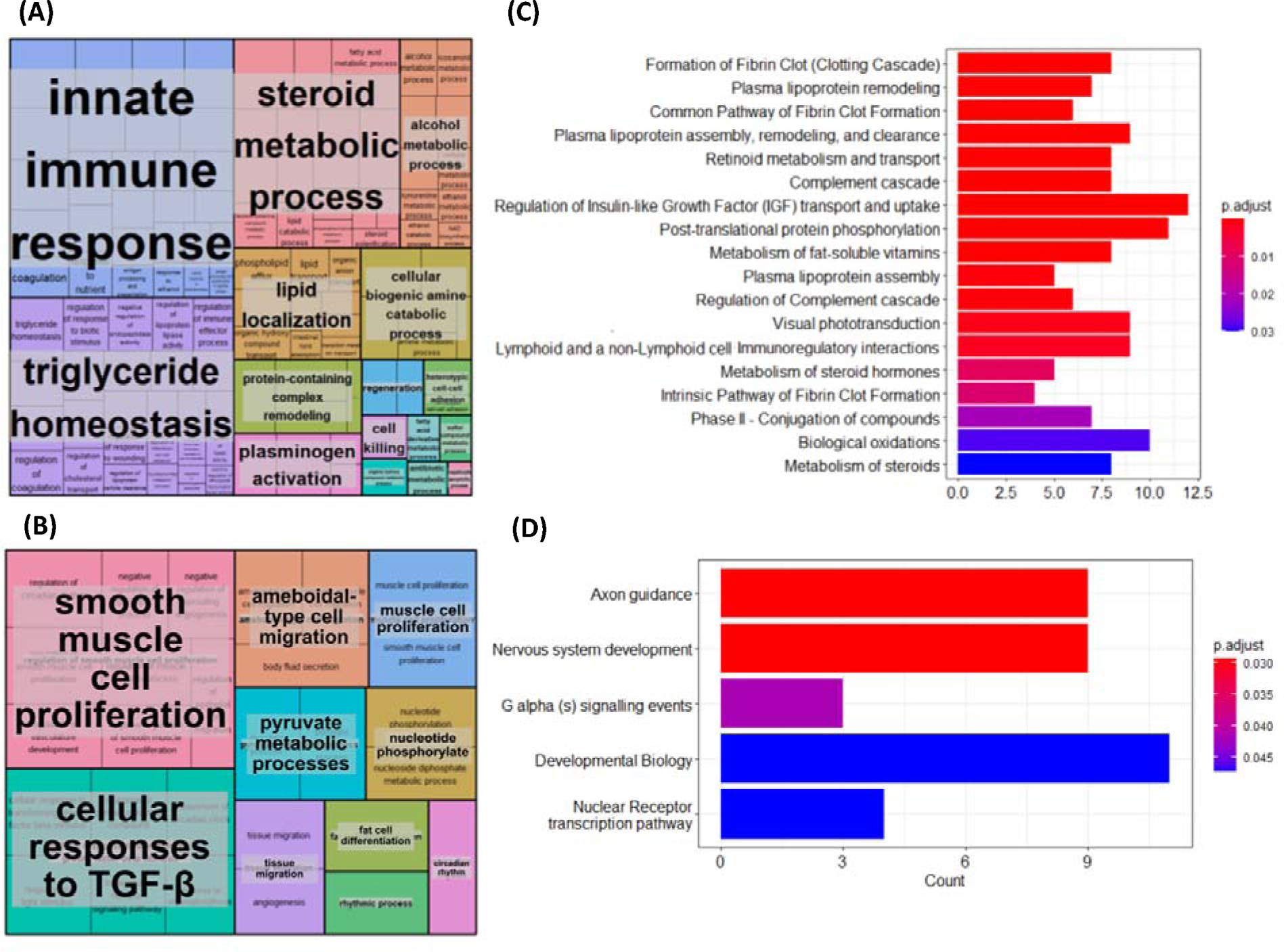
Gene Ontology (GO) and Reactome pathway analysis of genes differentially expressed in aged rat muscle but normalised to youthful expression by caloric restriction (CR). ClusterProfiler (version 3.14.3) package in R (version 4.1), was used to perform GO enrichment analysis. All the genes detected in skeletal muscle by RNA-seq analysis was used as background. A padj <0.05 was used as the cut off p-value. Revigo (http://revigo.irb.hr) then summarised GO terms for genes A. suppressed by and B. rescued by CR. Subsequently, the ReactomePA (version 3.15) package in R revealed Reactome pathways significantly (p-adj value=<0.05) associated with genes C. suppressed by and D. rescued by CR.

ClusterProfiler revealed 47 GO terms to be significantly associated with genes rescued in aged CR rats compared to aged rats fed AL. Top terms based on gene size included regulation of circadian rhythm, ATP generation from ADP and muscle cell proliferation. Revigo summarised genes suppressed by CR to primarily be involved in the regulation of smooth muscle cell proliferation, cellular response to transforming growth factor beta, ameboidal-type cell migration, and muscle cell proliferation (Figure 3B). Subsequently, genes suppressed by CR were found to be significantly enriched for 5 Reactome pathways, including axon guidance, nervous system development, G alpha (s) signalling events, developmental biology and nuclear receptor transcription pathway (Figure 3D).

### Functional and pathway analysis of protein-coding genes differentially expressed uniquely in CR rats

ClusterProfiler revealed 14 GO terms to be significantly associated with genes uniquely upregulated in age CR rats. Functional enrichment analysis showed that genes overexpressed under CR were linked to fatty acid metabolic process, white fat cell differentiation, carboxylic acid transport, organic acid transport and regulation of neurotransmitter levels, amongst others GO terms. Revigo summarised these 14 GO terms into 4 categories, these were: regulation of lipid metabolic process, fatty acid metabolic process, carboxylic acid transport and white fat cell differentiation (Figure 4C). As for pathways, we found one significant term associated with genes that were over-expressed under CR, which was biological oxidations.

**Figure 4.**
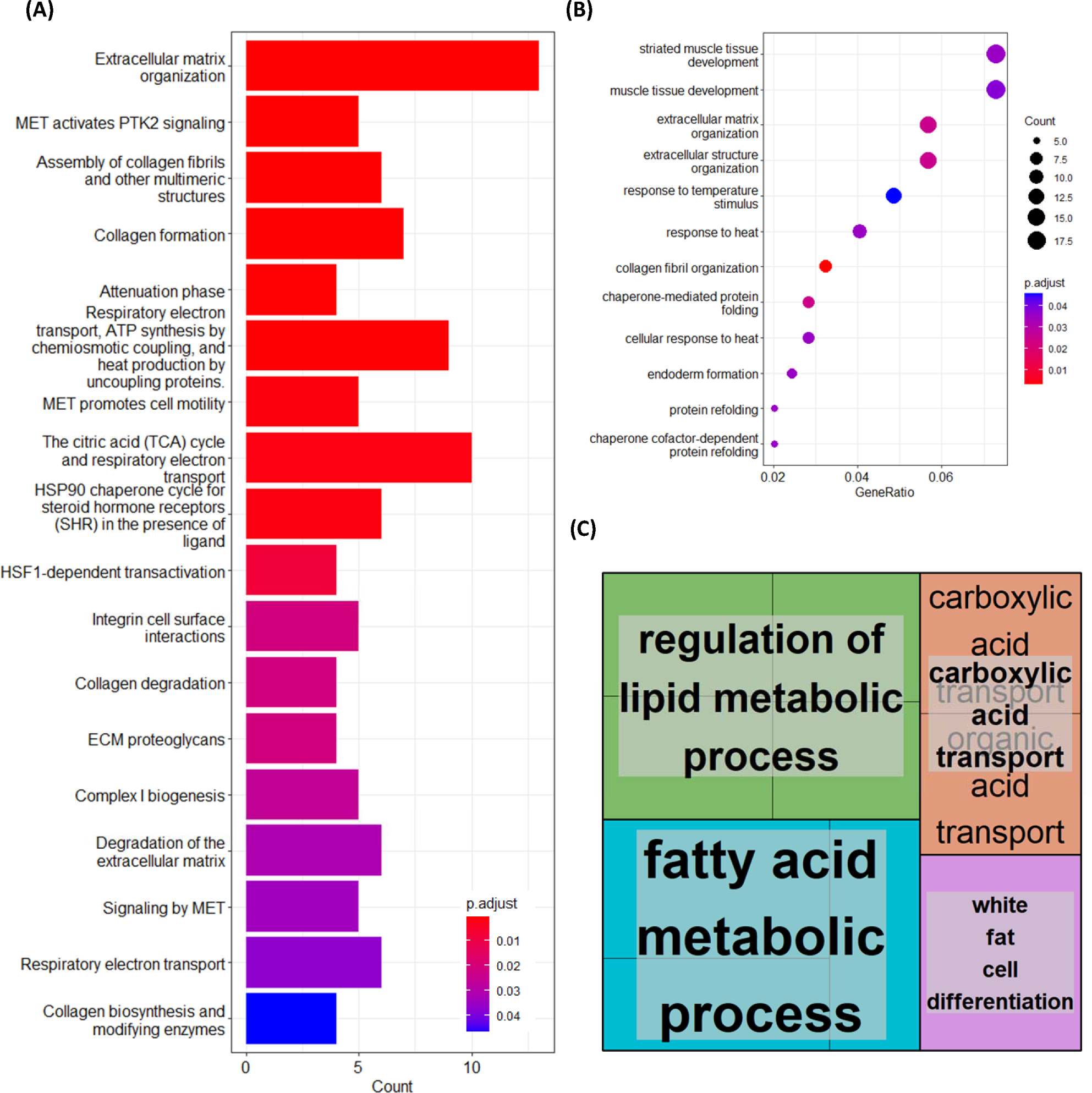
Gene Ontology (GO) and Reactome pathway analysis of genes differentially expressed in the muscle of aged caloric restriction (CR) but not aged non-CR rats. ClusterProfiler (version 3.14.3) and ReactomePA (version 3.15) packages in R (version 4.1) were used to perform GO and Reactome pathway enrichment analyses. All the genes detected in skeletal muscle by RNA-seq analysis was used as background. A padj <0.05 was used as the cut off p-value. For downregulated genes, significantly associated A. pathways and B. GO terms were revealed. Revigo could not summarise GO terms significantly associated with downregulated genes due to the low number of terms. C. For upregulated genes, GO terms were revealed, which Revigo (http://revigo.irb.hr) summarised. Only one pathway was found to be over-expressed in CR, which was biological oxidation.

ClusterProfiler revealed 12 GO terms to be significantly associated with genes uniquely downregulated by CR. Top terms based on gene size included muscle tissue development, extracellular matrix (ECM) organization, chaperone-mediated protein folding and collagen fibril organization (Figure 4B). Revigo clustering was not applied given the limited number of terms. Nonetheless, genes uniquely downregulated by CR were found to be significantly enriched for 18 Reactome pathways. These included ECM organization, MET activates PTK2 signalling, the citric acid (TCA) cycle and respiratory electron transport, integrin cell surface interactions and respiratory electron transport (Figure 4A).

### Functional and pathway analysis of protein-coding genes differentially expressed in ageing but unaffected by CR

Finally, we investigated the biological processes and the pathways associated with genes commonly upregulated in both the aged muscle of rats fed AL and CR when compared to young rats. ClusterProfiler revealed 254 GO terms to be significantly associated with genes commonly upregulated in aged muscle. Top terms based on gene size included lymphocyte-mediated immunity antigen processing and presentation of peptide antigen via MHC class I, cell killing, leukocyte-mediated immunity, acute inflammatory response and adaptive immune response regeneration (Figure 5A). Revigo clustering subsequently summarised these genes to be involved in the positive regulation of adaptive immune response based on somatic recombination of immune receptors built from immunoglobulin superfamily domains, the acute inflammatory response, long chain fatty acid metabolic processes, antigen processing and presenting and cell killing and regeneration. Subsequently, upregulated genes in aged rats were found to be significantly enriched for 21 Reactome pathways, including Platelet degranulation, Response to elevated platelet cytosolic Ca2+, Homeostasis and Regulation of Complement cascade.

**Figure 5.**
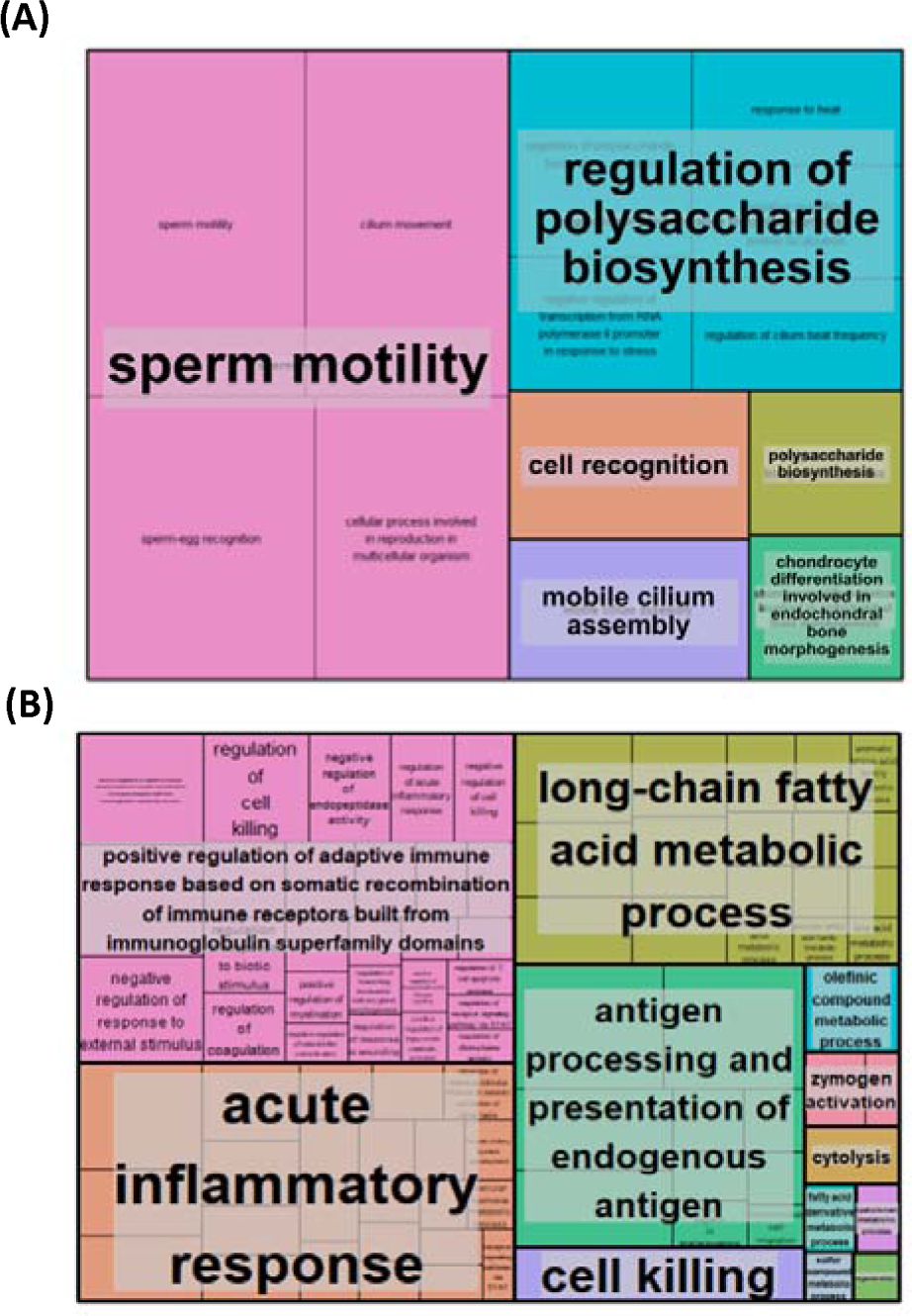
Gene Ontology (GO) analysis of genes differentially expressed in aged rat muscle but unaffected by caloric restriction (CR). ClusterProfiler (version 3.14.3) package in R (version 4.1), was used to perform GO enrichment analysis. All the genes detected in skeletal muscle by RNA-seq analysis was used as background. A padj <0.05 was used as the cut off p-value. Revigo (http://revigo.irb.hr) then summarised GO terms for A. upregulated genes and B. downregulated genes.

ClusterProfiler also revealed 33 GO terms to be significantly associated with genes commonly downregulated in aged muscle. The biological processes most commonly downregulated were found to include sperm motility, cellular processes involved in reproduction in multicellular organisms, cell-cell recognition, and regulation of glucan biosynthetic process. Revigo clustering subsequently summarised these genes to be involved primarily in sperm motility and the regulation of polysaccharide biosynthesis (Figure 5B). Pathway analysis demonstrated commonly downregulated genes to be associated with 3 significant Reactome pathways. These pathways were the HSP90 chaperone cycle for steroid hormone receptors (SHR) in the presence of ligand, attenuation phase and cellular response to heat stress.

### Weighted gene expression clustering analysis

Weighted Gene Correlation Network Analysis (WGCNA) revealed four co-expression modules, with two modules, turquoise and yellow, significantly correlated with ageing muscle in old rats. These modules were labelled as ageing-associated due to their relevance to ageing in old rats compared to young ones.

Further analysis focused on assessing the association between module membership (MM) and gene significance (GS) for ageing muscle. Genes within the turquoise and yellow modules were considered hub genes if they met stringent criteria (GS > 0.80 and MM > 0.80). Ultimately, two hub genes,

Enox1 and Slco1a4, were identified from the turquoise module, while 29 hub genes, including *Nlrc5, Bid*, *Nfkb2*, *Blm* and *Slc41a3*, were identified from the yellow module (Table 4). Among these hub genes, nine were upregulated and 22 were downregulated in ageing muscle compared to young muscle (Figure 6).

**Figure 6.**
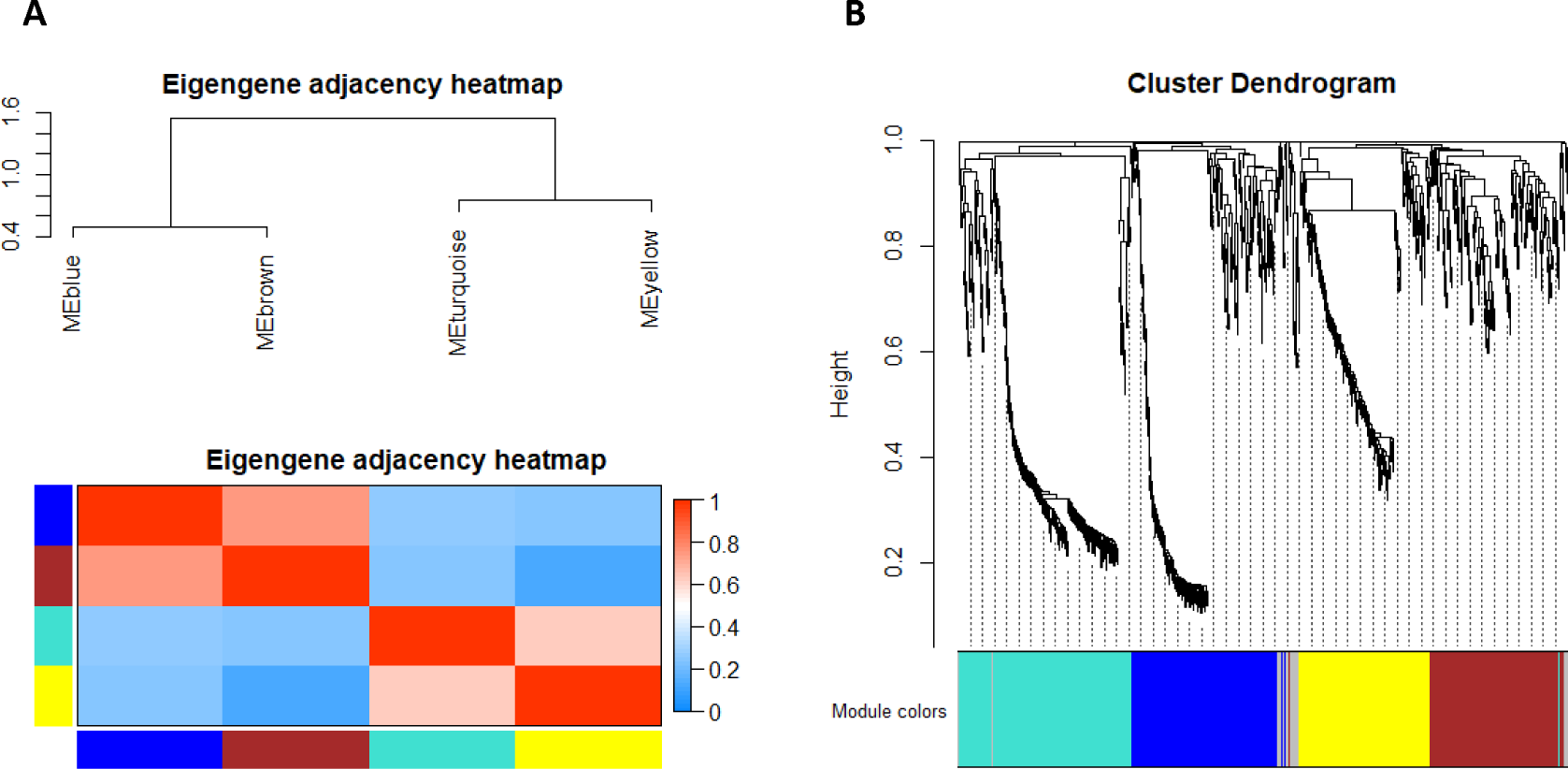
Weighted gene correlation network analysis (WGCNA) for DEG in old rat skeletal muscle. A. Module eigengene adjacency heatmap was clustered and were merged by a cut-off value of 0.25 and classified as the first principal component of a co-expression module matrix. Heatmap showing the similarity between 4 co-expressed modules (Blue, Brown, Turquoise, and Yellow). Colours indicate the strength of relation between the modules, red is highly associated, and blue shows no relation. B. Hierarchical clustering of the topological overlap matrix for DEG data between young vs old muscle. Colours at the bottom of the dendrogram denote various clusters, which is detected by the dynamic tree cut algorithm.

**Table 4.** Ageing-associated hub genes from turquoise and yellow modules. Weighted gene correlation network analysis revealed several hub genes, defined as any gene with a gene significance value >0.80 and module membership value >0.80. A. Shows hub genes from the turquoise module. B. Shows hub genes from the yellow module.

**A. Hub genes from turquoise module**
| Gene | log<sub>2</sub>(FoldChange) | p-value | padj |
| --- | --- | --- | --- |
| Enox1 | 0.95667696 | 1.16E-05 | 0.000842 |
| Slco1a4 | 1.055468763 | 1.52E-07 | 2.74E-05 |

| Gene | log<sub>2</sub>(FoldChange) | p-value | padj |
| --- | --- | --- | --- |
| Fcgr3a | 2.242886283 | 5.59E-07 | 7.82E-05 |
| Nlrc5 | 2.102821373 | 3.19E-12 | 3.35E-09 |
| RT1-A2 | 1.561580245 | 3.33E-16 | 1.12E-12 |
| Itgax | 1.454280188 | 2.84E-09 | 8.99E-07 |
| B2m | 1.266408587 | 4.57E-13 | 7.68E-10 |
| Bid | 1.163977519 | 2.76E-12 | 3.09E-09 |
| Negr1 | 1.134574724 | 1.43E-05 | 0.000994 |
| RT1-CE10 | 1.128927125 | 4.40E-09 | 1.34E-06 |
| Igtp | 1.009702659 | 8.94E-15 | 2.14E-11 |
| Adgre1 | 1.002848962 | 0.000356 | 0.011089 |
| Ephx1 | 0.978832197 | 2.14E-06 | 0.000225 |
| Chtf18 | 0.89566596 | 0.000397 | 0.011956 |
| Nfkb2 | 0.886183107 | 7.68E-10 | 2.93E-07 |
| Stat4 | 0.730178567 | 2.23E-07 | 3.46E-05 |
| Blm | 0.703401124 | 5.63E-05 | 0.002862 |
| Nme1 | 0.696274908 | 2.41E-05 | 0.001445 |
| Lrrc14 | 0.65053891 | 9.65E-05 | 0.004389 |
| Ap4m1 | 0.623698112 | 0.000136 | 0.005508 |
| Rcn1 | 0.598694992 | 5.36E-07 | 7.62E-05 |
| Ppia | 0.593784395 | 4.84E-07 | 7.13E-05 |
| Synpo | -0.695219835 | 0.000189 | 0.007002 |
| Grb10 | -0.707694024 | 0.000122 | 0.00507 |
| Rilpl1 | -0.734518172 | 0.000463 | 0.013262 |
| Cry1 | -0.752452225 | 7.20E-08 | 1.47E-05 |
| Mb | -0.761868075 | 0.000302 | 0.009895 |
| Ubc | -0.877083539 | 6.59E-05 | 0.003233 |
| Dnajb5 | -0.938064658 | 8.59E-06 | 0.000656 |
| Asb2 | -0.960869149 | 4.64E-05 | 0.002493 |
| Slc41a3 | -1.004164203 | 1.01E-06 | 0.00013 |

Using the same approach as that used to identify the ageing hub genes, WGCNA identified one hub gene, *Zfp950*, from the yellow module and 20 hub genes from the blue module. The combined hub gene set from both modules is presented in a table (Table 5). Examining the impact of muscle ageing under CR, we observed that among the identified hub genes, five were downregulated, including *Arpp21, LOC100362814, Prr32, Sipa1l2* and *Tnfrsf12a*. Meanwhile, 16 genes, including *Blm, Cep112, Fam214a* and *Zfp950*, were upregulated. These findings shed light on specific genes affected by the ageing process under calorie restriction in muscle tissue (Figure 7).

**Figure 7.**
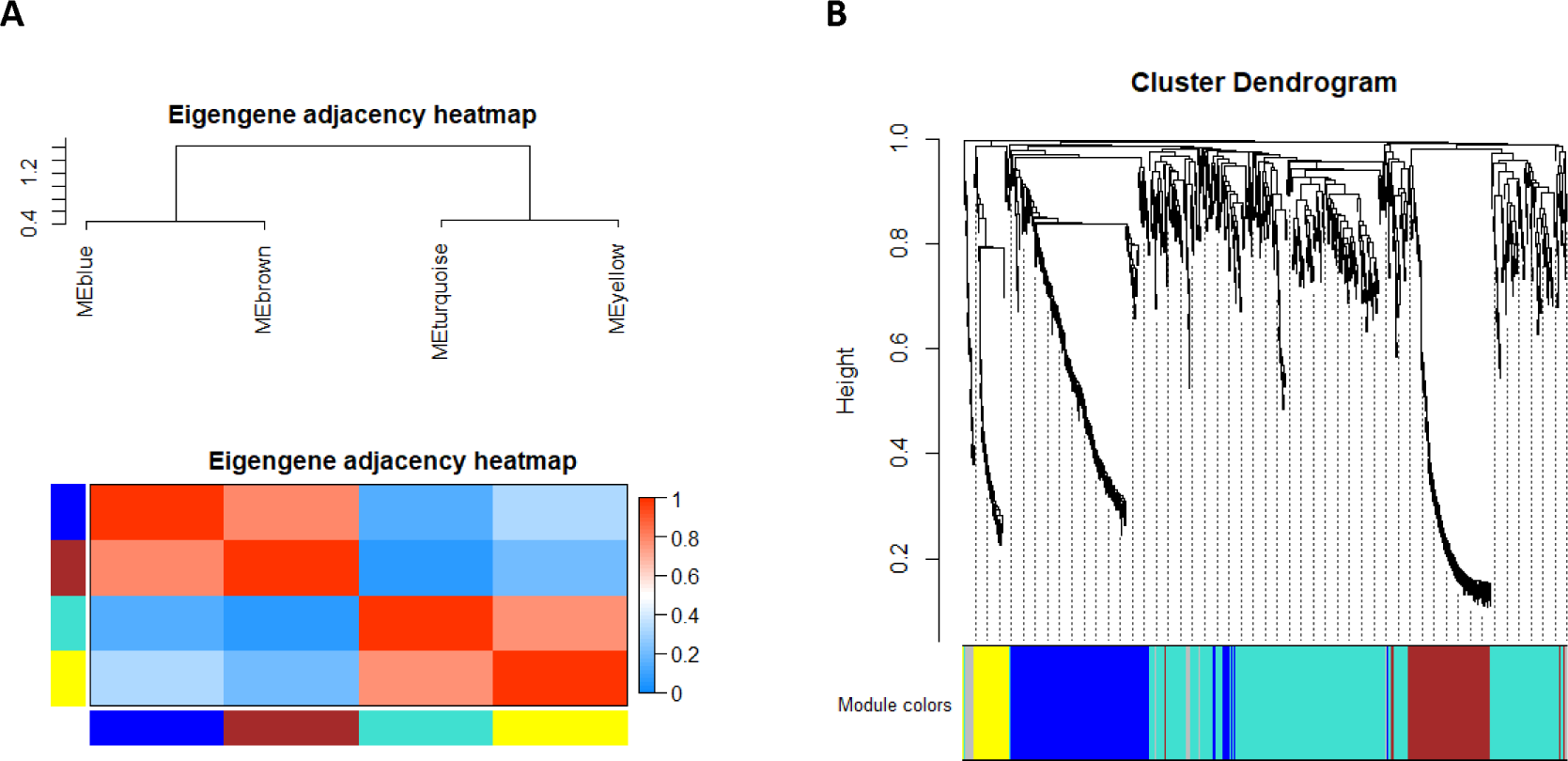
Weighted gene correlation network analysis (WGCNA) of DEG in old CR skeletal muscle from rat. A. Module eigengene adjacency heatmap was clustered and were merged by a cut-off value of 0.25 and classified as the first principal component of a co-expression module matrix. Heatmap showing the similarity between 4 co-expressed modules (Blue, Brown, Turquoise, and Yellow). Colours indicate the strength of relation between the modules, red is highly associated, and blue shows no relation. B. Hierarchical clustering of the topological overlap matrix for DEG data between young vs old CR muscle. Colours at the bottom of the dendrogram denote various clusters, which is detected by the dynamic tree cut algorithm.

**Table 5.** CR-associated hub genes from blue and yellow modules. Weighted gene correlation network analysis revealed several hub genes, defined as any gene with a gene significance value >0.80 and module membership value >0.80.

| Gene | log<sub>2</sub>(FoldChange) | pvalue | padj |
| --- | --- | --- | --- |
| Arpp21 | -1.345552593 | 1.23E-08 | 2.19E-06 |
| Blm | 0.770525928 | 4.17E-08 | 6.20E-06 |
| Cep112 | 0.692489655 | 5.64E-05 | 0.00228 |
| Chaf1a | 0.765123713 | 6.89E-06 | 0.000481 |
| Fam214a | 0.841061048 | 3.39E-07 | 3.70E-05 |
| Fmo1 | 1.334894874 | 6.96E-10 | 1.85E-07 |
| Glul | 0.700178742 | 2.81E-06 | 0.00023 |
| LOC100362814 | -0.609744488 | 0.000106 | 0.003563 |
| Meis1 | 0.612209741 | 0.000109 | 0.003652 |
| Mthfd1l | 0.86672007 | 2.74E-05 | 0.001299 |
| Prr32 | -0.779158274 | 1.97E-05 | 0.001017 |
| RGD1305184 | 2.595063618 | 5.12E-07 | 5.36E-05 |
| RGD1566386 | 0.591300765 | 7.38E-05 | 0.002769 |
| RT1-A1 | 0.655507604 | 6.16E-06 | 0.000439 |
| RT1-A2 | 1.058148782 | 8.29E-10 | 2.06E-07 |
| RT1-S3 | 1.008016385 | 4.52E-07 | 4.82E-05 |
| Scin | 1.13040547 | 2.40E-05 | 0.001183 |
| Sipa1l2 | -0.993686971 | 3.03E-10 | 8.93E-08 |
| Tnfrsf12a | -0.871819879 | 5.00E-06 | 0.000367 |
| Vrk1 | 0.600482485 | 2.46E-06 | 0.00021 |
| Zfp950 | 0.626220825 | 1.01E-07 | 1.31E-05 |

### PPI network analysis of ageing-associated and CR-associated differentially expressed genes in aged muscle

To explore ageing-related genes in muscle, PPIs were investigated using the STRING database, which facilitated the identification of known and predicted interaction between proteins for the genes of interest. The yellow module exhibited 172 nodes and 181 edges in its interaction networks, while the turquoise module contained 232 nodes and 665 edges. Key proteins within these networks were identified. In the yellow module, four proteins—*Ifit3, Irf7, Rsad2* and *Usp18*—stood out across different methods, suggesting their significance in muscle ageing (Figure 8, Table 6A). These proteins were labelled as ageing-related hub genes, all showing upregulation compared to younger samples.

**Figure 8.**
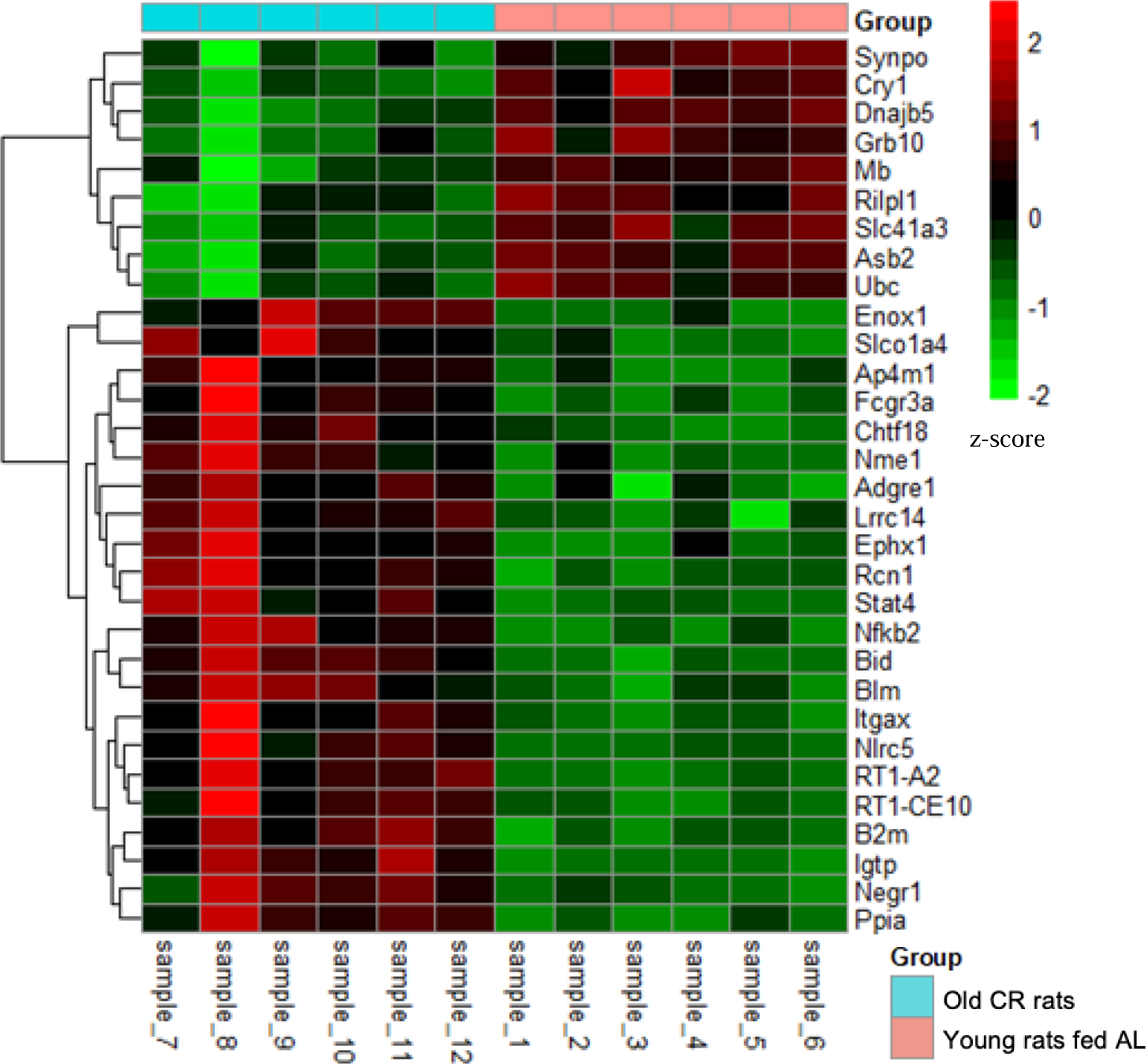
Heatmaps represent the expression level of hub genes from turquoise and yellow module. Pheatmap package (version 1.0.12) was used to create the heatmap. Normalised read counts of hub genes were used and z-score were computed to produce the heatmap. Red colours represent higher expression and green represent lower expression. X-axis showing samples and y-axis showing genes. Group colour turquoise represents old skeletal muscle and pink represents young skeletal muscle of.

**Table 6.**
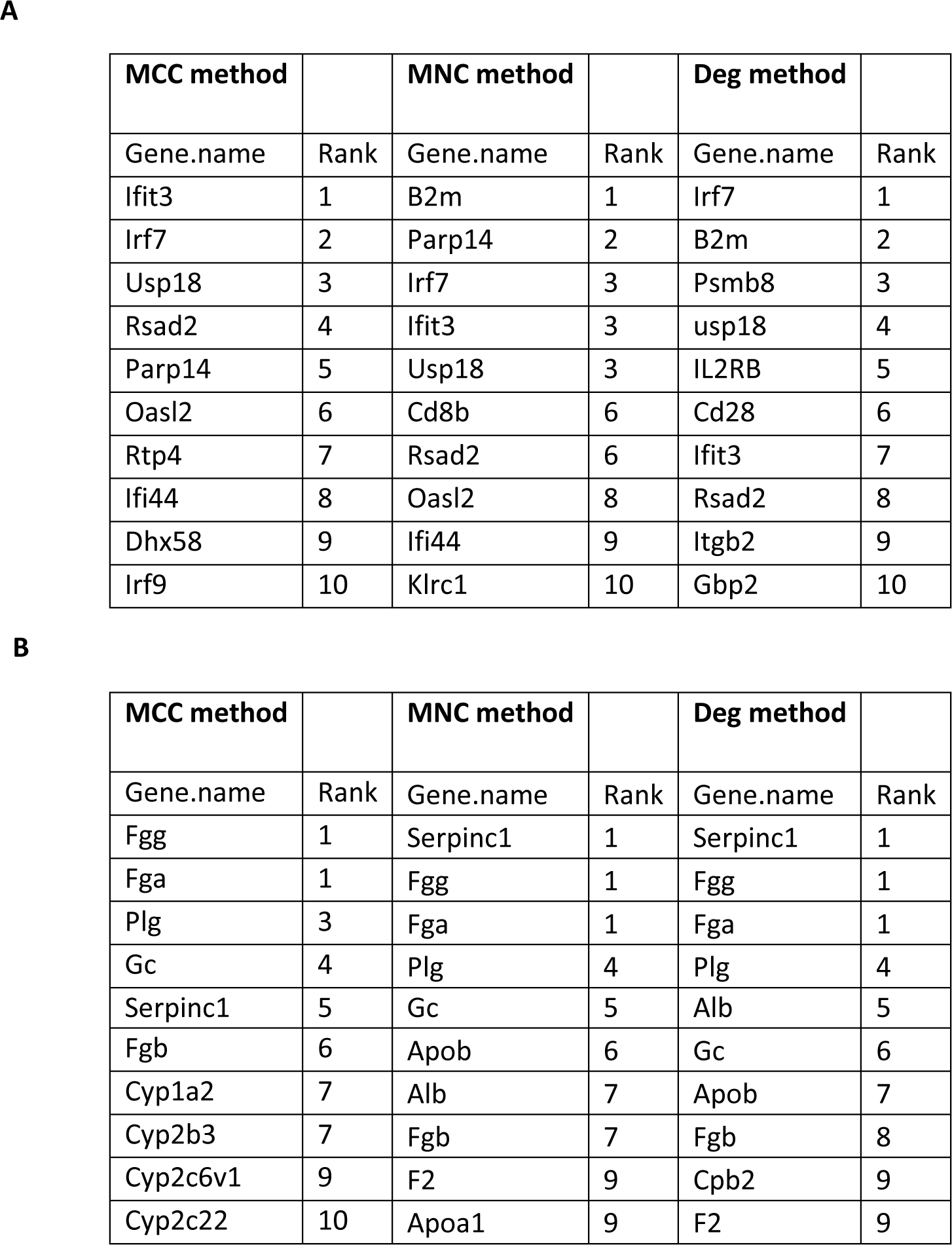
Ageing-associated protein hubs in rat muscle. Results from CytoHubba analysis for (A) the yellow module and (B) the turquoise module. CytoHubba plugin of Cytoscape (version 3.9.1) was used to rank the nodes by their network features. Top 10 ranked nodes were obtained using MCC, MNC and Deg method from CytoHubba.

Interestingly, *B2M*, a gene well-characterised to display age-associated dyregulation, appeared both in the PPI network analysis and the WGCNA method. In the turquoise module, six proteins—*Fgg, Fga, Plg, Gc, Serpinc1* and *Fgb*—emerged as prominent hub genes associated with ageing (Figure 8, Table 6B). Notably, all six showed upregulation in ageing muscle. However, when comparing these hub genes to the WGCNA method, no overlapping genes were found, indicating distinct findings.

To explore genes associated with muscle ageing in CR rats, PPIs were investigated using the STRING database. The yellow module exhibited 51 nodes with 72 edges in its interaction network, while the blue module contained 224 nodes and 157 edges. Key proteins within these networks were identified. In the yellow module, eight hub genes emerged as significant across the methods—*GC, Alb, Apoa1, Ambp, Plg, F2, Apoh* and *Orm1* (Figure 9, Table 7A). Similarly, all these hub genes displayed upregulation in the aged muscle of CR rats when compared to the aged muscle of rats fed AL. In the blue module, seven proteins—*Irf7, Ifit3, Usp18, Mx1, Rsad2*, *Oasl2* and *Rtp4*—were significant across all three methods (Figure 9, Table 7B). These proteins all exhibited upregulation in the aged muscle of CR rats when compared to the aged muscle of rats fed AL. Notably, however, no common hub genes were found between the CytoHubba methods and the WGCNA (MM vs GS).

**Figure 9.**
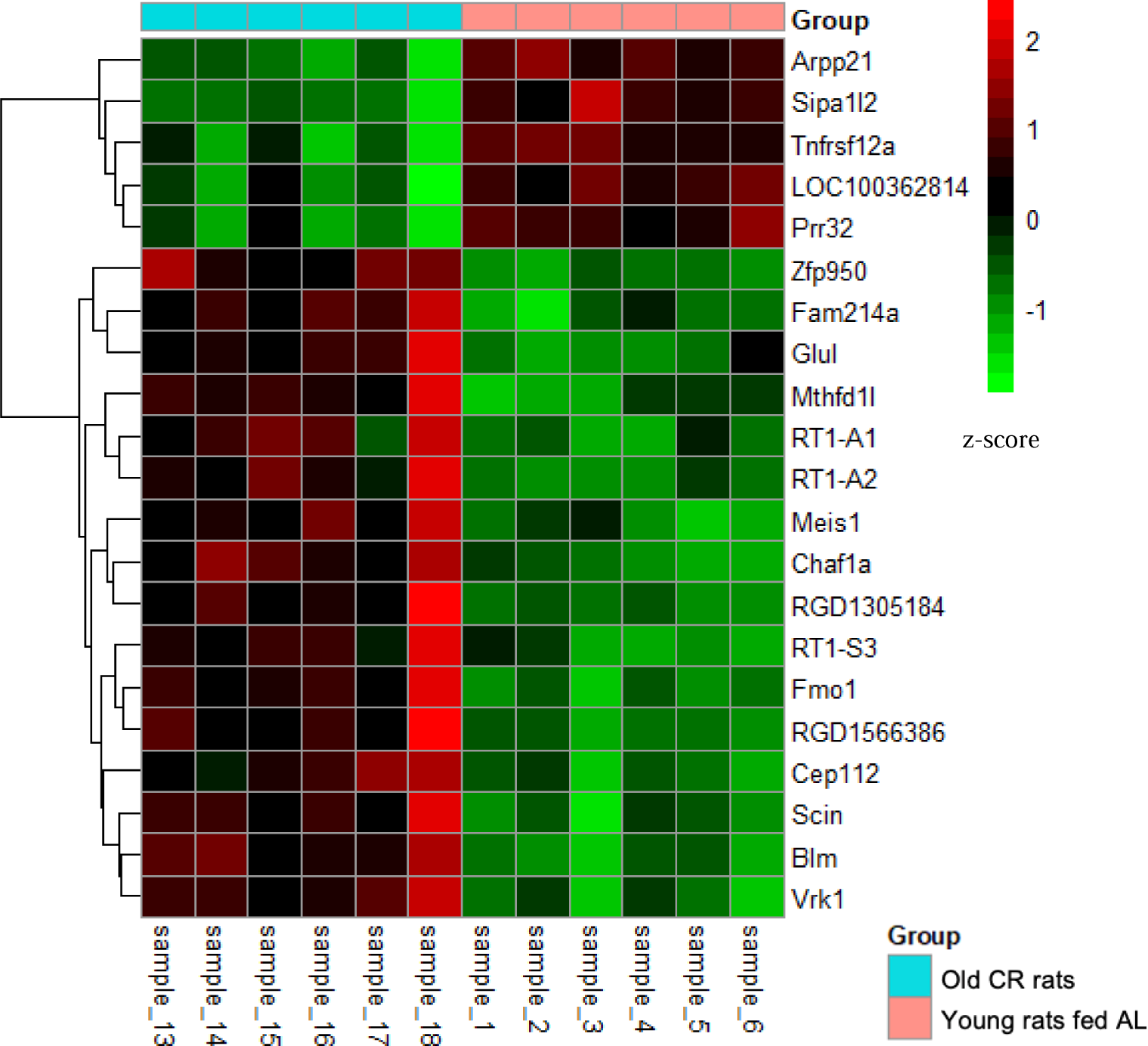
Heatmaps represent the expression level of hub genes from yellow and blue module. Pheatmap package (version 1.0.12) was used to create the heatmap. Red colours represent higher expression and green represent lower expression. X-axis showing samples and y-axis showing genes. Group colour turquoise represents old CR skeletal muscle and pink represents young skeletal muscle of rat. Sample 1-6 represent 6-month-old and sample 13-18 represent 28 months CR.

**Table 7.**
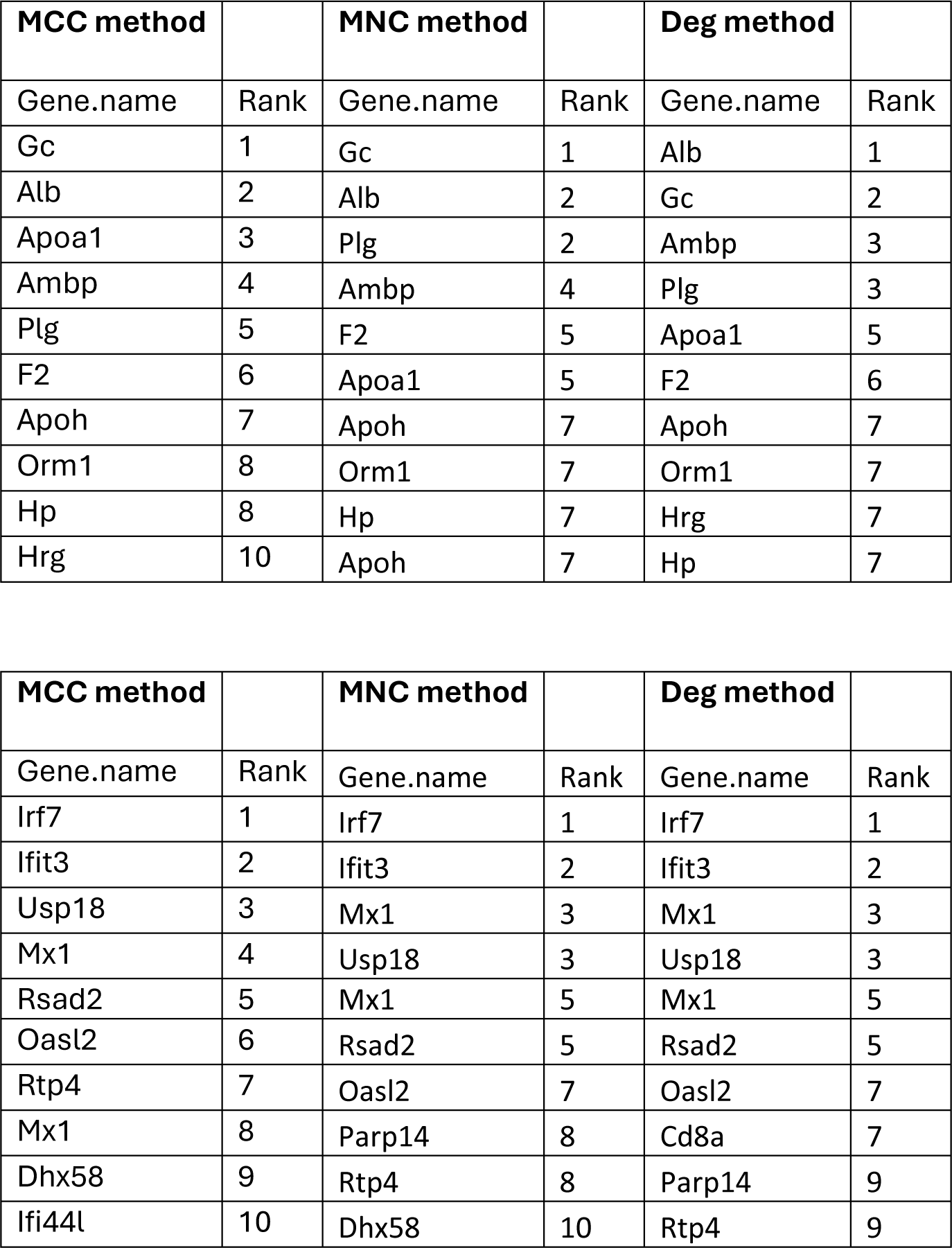
CR-associated protein hubs in aged rat muscle. Results from CytoHubba analysis for (A) the yellow module and (B) the blue module. CytoHubba plugin of Cytoscape (version 3.9.1) was used to rank the nodes by their network features. Top 10 ranked nodes were obtained using MCC, MNC and Deg method from CytoHubba.

### Analysis of overlap between Ageing-associated and CR-associated hub genes

The WGCNA method found only 2 hub genes to be affected by both ageing and CR, *Blm* and *RT1-A2*. However, PPI methods identified 6 hub genes to be affected by both ageing and CR (Figure 10, Table 8). These included *Gc, Plg, Irf7, Ifit3, Usp18* and *Rsad2*. Nonetheless, 13 out of 19 hub genes associated with either ageing or CR were unique to one or the other. Hub genes unique to old CR muscle included *Alb, Apoa1, Ambp, F2, Apoh, Orm1, Mx1, Oasl2* and *Rtp4*, meanwhile, hub genes unique to old muscle included *Fgg, Fga, Serpinc1* and *Fgb*.

**Figure 10.**
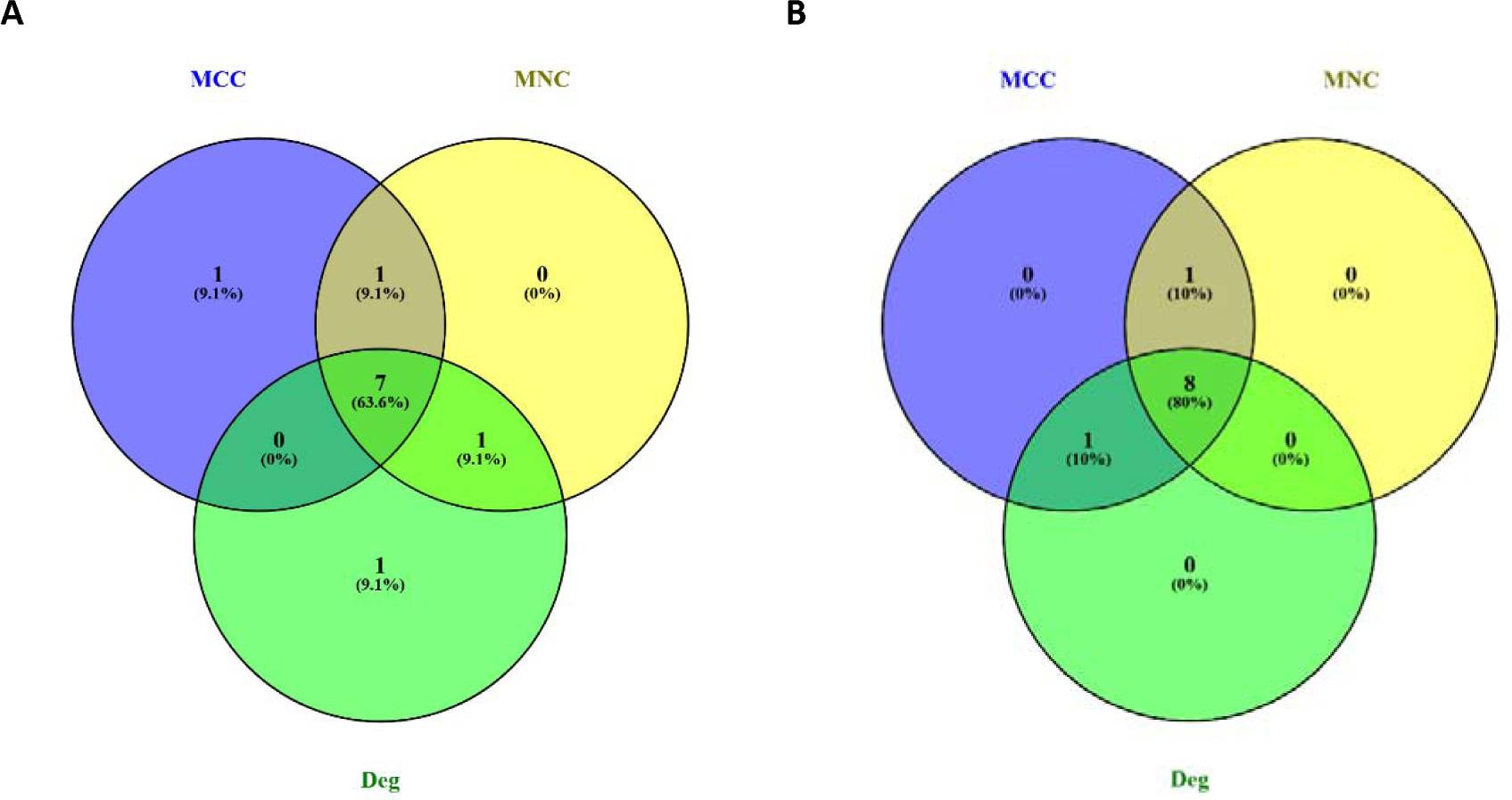
Venn diagrams showing the degree of overlap between results using various method to identify hub protein involved in the effects of A. ageing and B. caloric restriction (CR) in rat skeletal muscle. Venn diagram was created using “ https://bioinfogp.cnb.csic.es/tools/venny. Hub genes were identified using cytoHubba via 3 different methods: Maximal Clique Centrality (MCC), Maximum Neighbourhood Component (MNC) and the Degree method (Deg). Hub genes identified by more than one method were assumed to be the least likely to be false-positive.

**Table 8.**
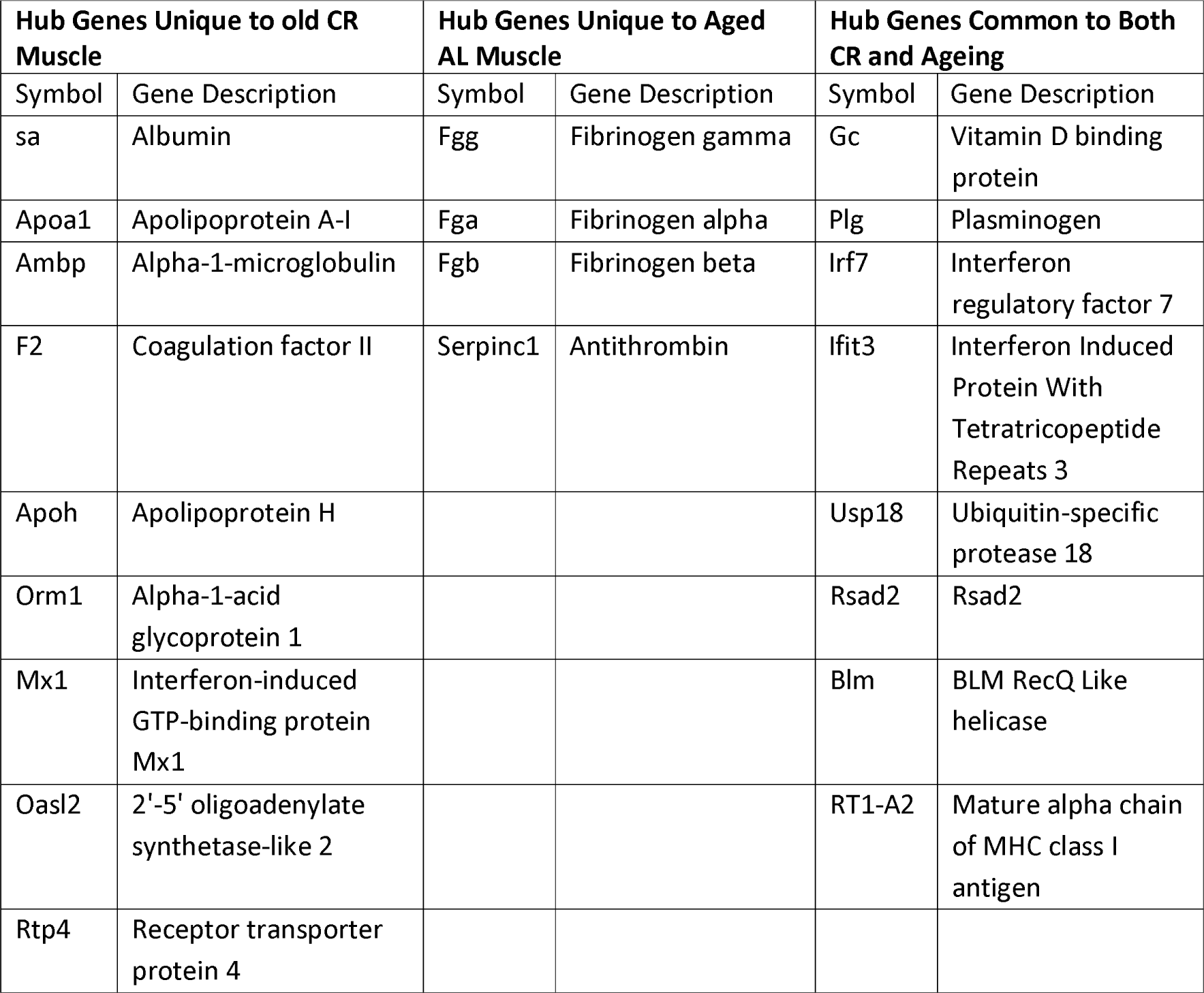
The overlap of hub genes identified by protein-protein interaction PPI network methods between the muscle of aged rats fed ad libitum and aged rats caloric restricted (CR). Eight out of twelve hub genes involved in ageing were also involved in responses to CR. This suggests an incomplete reversal of the ageing phenotype by CR. Nine hub genes were found to mediate the unique effects of CR, independent of ageing reversal.

### Validation of RNA-seq analysis using qRT-PCR

RNA-seq results suggested all 6 of the genes selected for validation to be significantly downregulated in the aged AL-fed rats when compared to the young AL-fed rats, and to be significantly upregulated in the aged CR rats when compared to the aged, AL-fed rats. On the other hand, RT-qPCR found only 4 out of 6 to be significantly downregulated [(*FOXO1*, p= 0.002), (*KLF4*, p= 0.02), (*CRY1*, p= 0.001), (*NR4A3*, p=0.004)] in the aged, AL-fed rats when compared with young AL-fed rats (Figure 11). *OGDH* and *PGC1A* were not significantly downregulated in old samples according to the RT-

**Figure 11.**
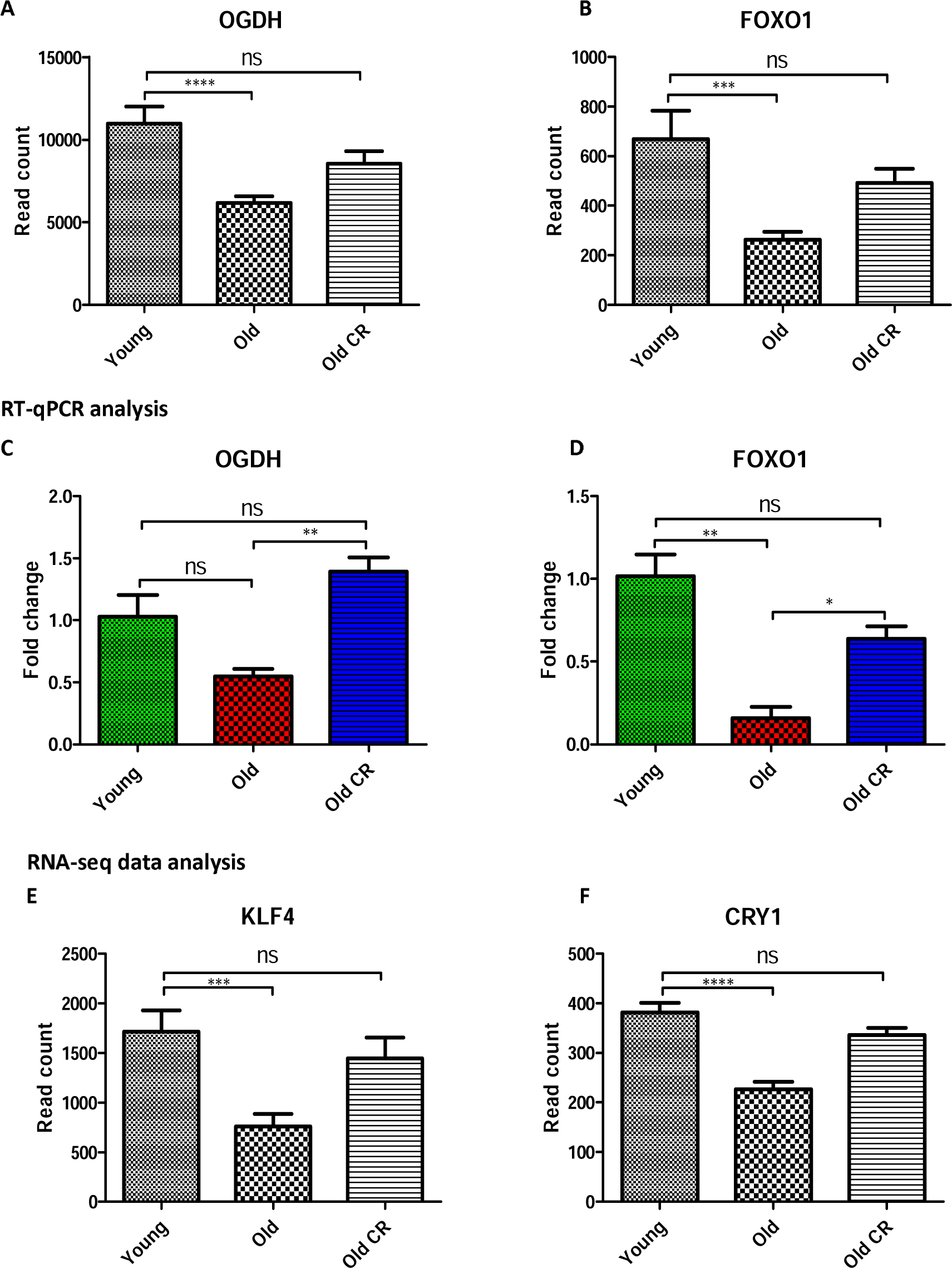
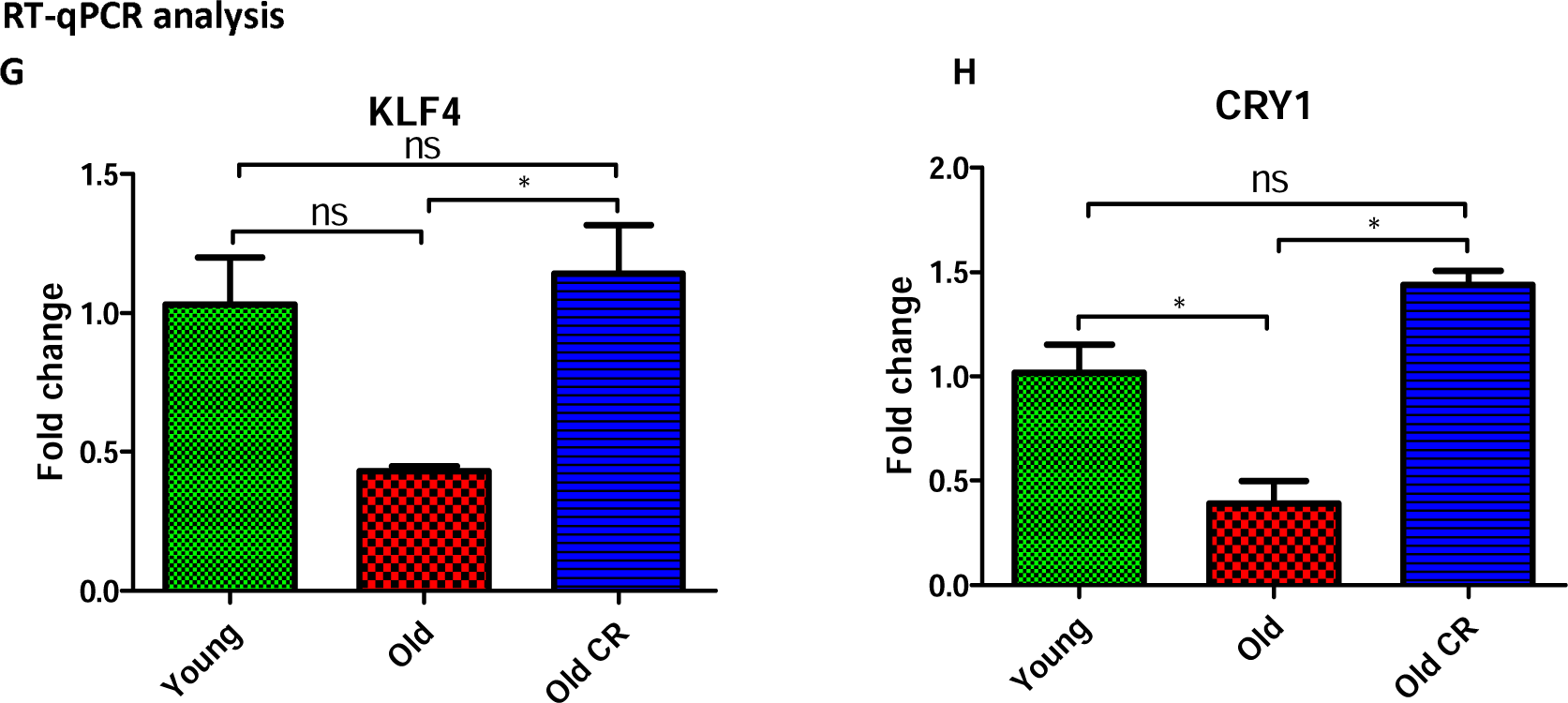
Normalised read count from RNA-seq data compared to relative expression levels in fold change from RT-QPCR. For rt-qpcr samples were calculated relative to Fbxo45 mRNA levels and were compared using One-Way ANOVA test. Of the 6 RNAs selected for analysis, statistically significant changes in expression with old and old-CR samples compared to young were identified for OGDH, FOXO1, KLF4, CRY1, PGC1A and NR4A3 mRNAs (A-L), p-values denoted by a star asterisk. All p-values are unadjusted for rt-qpcr. Error bars represent mean1±1SEM.

qPCR analysis, potentially due to limited sample size (n=3). Expression of *OGDH* was higher in the aged rats fed AL, but not significantly so (Figure 12). Nonetheless, consistent with RNA-seq results, RT-qPCR analysis suggested all 6 selected genes to be upregulated/rescued in the aged CR rats when compared to the aged, AL-fed rats.

**Figure 12.**
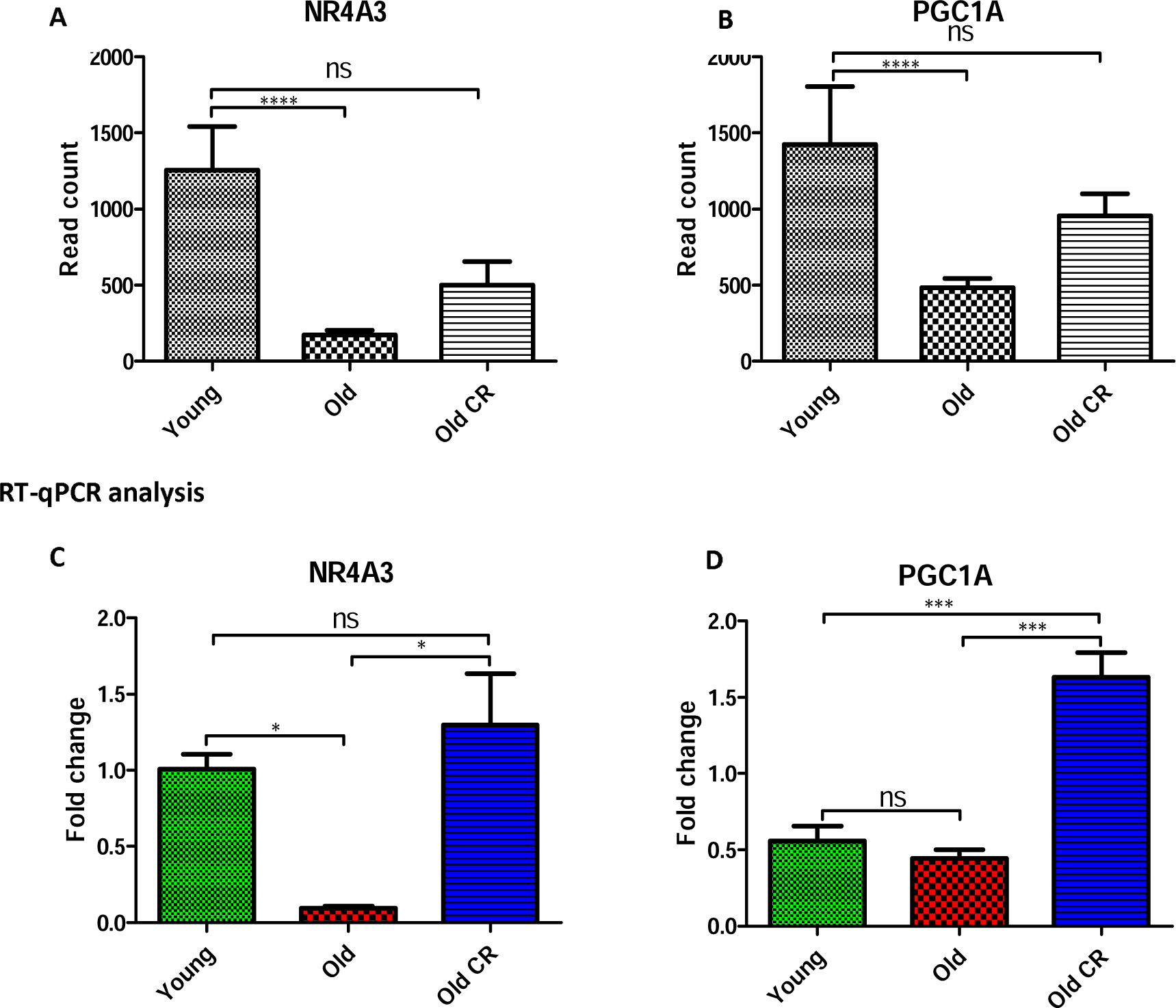
Normalised read count from RNA-seq data and relative expression levels in fold change from RT-QPCR. For rt-qpcr samples were calculated relative to Fbxo45 mRNA levels and were compared using One-Way ANOVA test. Of the 6 RNAs selected for analysis, statistically significant changes in expression with old and old-CR samples compared to young were identified for OGDH, FOXO1, KLF4, CRY1, PGC1A and NR4A3 mRNAs (A-L), p-values denoted by a star asterisk. All p-values are unadjusted for rt-qpcr. Error bars represent mean1±1SEM.

## Discussion

### Differentially Expressed Genes

As expected, we found many genes to be differentially expressed between the muscle of aged and youthful rats fed AL. Furthermore, consistent with previous findings from both rats, mice and humans ^11,34^, CR was found to suppress 69.7% and rescue 57.8% of the genes found to be upregulated and downregulated in aged muscle, respectively. Our previous study found CR to reverse differential expression of a similar percentage of transcripts in the brain of these animals ^35^. CR reversed a majority of the age-related changes in transcriptomic profiles.

GO and pathway enrichment analysis of differentially expressed genes replicated the well-characterised finding that ageing regardless of tissue type is generally characterised by immune/stress response gene over-expression and metabolic/developmental gene under-expression ^36^. Indeed, the genes most significantly upregulated in aged rats fed AL were associated with protein folding and immune responses, corroborating influential theories of ageing ^37,38^. CR suppressed many of the genes associated with these processes, normalising the aged phenotype. Interestingly, consistent with programmatic accounts of ageing ^39^, the genes that were most significantly downregulated in aged rats fed AL were genes related to developmental biology. These genes were rescued by CR.

While CR did normalise many biological pathways perturbed in ageing, it also uniquely altered pathways that are perhaps crucial for normal muscle function. Genes upregulated uniquely by CR in aged rats were most associated with fatty acid metabolism, meanwhile, those downregulated uniquely by CR rats were most associated with ECM organisation and cellular respiration. Although several of these changes are likely involved in the metabolic shift towards increased fat utilisation with CR ^40^, effects on the ECM may contribute to unwanted side-effects. Indeed, reduced ECM synthesis is thought to contribute to exacerbated age-associated deficits in wound healing ^41^ and skin quality ^42^ concomitant with ageing. Saying this, reducing ECM deposition has been shown to contribute to the anti-hypertensive ^43^ and neuroprotective ^44^ effects of CR. Ultimately, how CR-mediated alterations in ECM organisation might alter sarcopenia and overall muscle health should be explored further.

### Ageing- and CR-associated Hub Genes

Several hub genes associated with both ageing and CR responses in rat muscle were identified. The function of these genes and how these might contribute to muscle ageing are listed below.

1. ***Blm*** (a 3′−5′ ATP-dependent RecQ DNA helicase) is a necessary genome stabilizer, which controls DNA replication, recombination and repair. Its dysregulation may perturb DNA repair processes within muscle cells, contributing to the accumulation of DNA damage with age ^45^.
2. ***RT1-A2*** (RT1 class Ia, locus A, sub-locus 2) is involved in presenting antigens to immune cells ^46^. Its dysregulation with ageing may contribute to chronic inflammation but its overall role in the ageing process is largely unexplored.
3. ***Gc*** (Vitamin D-binding protein) is responsible for the transport and regulation of vitamin D precursors and metabolites, implicating dysregulated Vitamin D signalling in age-related muscle decline, as has been shown previously ^47^.
4. ***Plg*** (Plasminogen) is a precursor of the enzyme plasmin, involved in breaking down blood clots. In muscle ageing, its dysregulation could impact repair processes and regeneration efficiency, as has been hypothesised previously ^48^.
5. ***Irf7*** (Interferon regulatory factor 7) is involved in interferon-mediated immune responses. Its overexpression (although only in adipose tissue) has been found to dysregulate mitochondrial function and amino acid metabolism ^49^. Furthermore, *Irf7* has recently been shown to be involved in the modulation of satellite cell proliferation and regeneration with age ^50^. These functions possibly contribute to its effects on muscle loss.
6. ***Ifit3*** (Interferon-induced protein with tetratricopeptide repeats 3) and ***Usp18*** (Ubiquitin-specific protease 18) are also involved in interferon-mediated immune responses. Their dysregulation with ageing may contribute to chronic inflammation phenotypes ^51,52^. *Usp18* is also a known regulator of muscle cell differentiation ^52^.
7. ***Rsad2*** (Radical S-adenosyl methionine domain-containing protein 2) plays a role in defence against viruses. Its involvement in muscle ageing likely relates to its ability to modulate cellular responses to stress, as has been shown in age-related macular degeneration ^53,54^ Furthermore, *Bid* and *B2m* also appeared to be involved both in ageing and in responses to CR. Although neither gene was identified as a CR-related hub gene, they were both strongly upregulated by CR and shown to be ageing-associated hubs genes. *Bid* (BH3 interacting domain death agonist), a pro-apoptotic gene ^55^, potentially contributes to muscle ageing by accelerating the apoptosis of differentiated and satellite cells. In addition to pro-apoptotic functions, *Bid* has been shown to be involved in metabolism and DNA damage ^56^. Conversely, *B2m* (β2-microglobulin), is linked to senescence ^57^, hinting at a potential role for senescence suppression in CR-mediated muscle ageing protraction. Additionally, *NFkB2* (Nuclear Factor Kappa B Subunit 2) was an ageing-associated hub gene, which is a well-characterised inflammatory regulator and known to be associated with the loss of muscle mass ^58^.

Some interesting hub genes uniquely associated with CR, included *ARPP21, VRK1, Glul* and *Orm1*. *ARPP21* is associated with myogenic differentiation ^59,60^. *VRK1* upregulation by CR, potentially affects muscle health positively via downstream AMPK activation ^61^. Lastly, dietary intake of the amino acid glutamine is thought to aid in the growth of maintenance of muscle mass but not in the elderly ^62^. It is interesting, therefore, that *Glul* (glutamine ammonia ligase) was overexpressed in the muscle of CR rats. *Orm1* (orosomucoid 1), a crucial acute-phase plasma protein, is involved in controlling angiogenesis and regulating inflammation and the immune system ^63^. However, its upregulation has also been further shown to enhance glycogen storage in the muscle, as well as increase muscle endurance ^64^. This finding suggests CR may also modulate genes that are involve in improving muscle strength and endurance via mechanisms independent of those affected by ageing.

### Limitations

The primary limitations of this study were the small number of biological replicates (n=6) and RNAs and the fact that they were collected from skeletal muscle stored over 18 years ago. However, the RNA samples prior to sequencing were assessed and passed the quality and integrity checks. We employed moderate criteria to screen the genes, in the data cleaning process and most importantly differentially expressed genes were partially validated by randomly selecting genes and samples, using qPCR.

Unfortunately, comparisons with other studies revealed limited overlap between the genes shown to be differentially expressed in aged muscle in our study and those reported in the literature. For example, only 10 genes were differentially expressed in aged muscle in both our study and another rat study ^14^, only 16 genes were differentially expressed in aged muscle in both our study and a human study ^65^ and only 16 genes were differentially expressed in aged muscle in both our study and a mouse study ^12^. There was however, a great degree of overlap (187 genes) between the genes identified by our study of rat muscle ageing and a study of mouse muscle ageing ^11^. Equally, our results displayed a good degree of overlaps (61 genes) of the muscle data in a meta-analysis of tissues-specific transcriptomic correlates of ageing ^36^.

Despite the limited overlap, genes consistently dysregulated in ageing muscle across various studies, such as *Cepb1*, are likely most strongly involved in muscle ageing and potentially in the development of sarcopenia. Nonetheless, the lack of replication suggests that more extensive investigations involving larger cohorts and diverse populations are necessary to elucidate common versus specific transcriptional changes in ageing muscle. Among the 903 genes differentially expressed in the aged muscle of CR rats when compared to the muscle of aged rats fed AL, 314 were also noted in prior studies across different species, indicating somewhat greater consistency in findings.

## Conclusion

Overall, using RNA-seq data from the muscle of young AL-fed and CR rats and aged AL-fed and CR rats, we identified changes in the of expression protein-coding genes consequent to both ageing and CR. Several of the identified genes were previously known to be involved in muscle ageing and responses to CR, replicating previous findings. Indeed, we found inflammatory genes to display a large degree of expression change in muscle ageing and this to be ameliorated by CR. In addition to known genes, our results implied genes involved in circadian rhythm regulation and developmental biology to perhaps play a more important role in muscle ageing than previously realised. Additionally, using co-expression analysis and PPI network analysis, we identified the key hub genes regulating these pathways. Knowledge about these key proteins may offer novel insights into the mechanisms via which the muscle ageing process could perhaps be perturbed, without having to implement CR. These key genes should be studied further in in-vitro or in-vivo ageing models. In summary our work indicates that CR is slowing down, but not completely preventing, age-related muscle changes. Some age-related muscle changes are still occurring even with CR, suggesting there are factors driving these changes that are not modified by CR feeding alone. More research is needed to further unravel these mechanisms. However, the partial protective effect of CR demonstrates the modifiable nature of the muscle ageing processes.

## Authors’ contributions

GA and JPM conceived the study. BJM provided materials. GA performed all the investigations and analysis. GA wrote the manuscript, with CWB structuring the manuscript. The final version of the manuscript was edited and revised by GA, CWB, JPM. The study was supervised by JPM, KGW. PR assisted with both computational and laboratory investigation. CWB assisted with bioinformatics analysis.

## Acknowledgements

We gratefully acknowledge the funding provided by CIMA for this project. Additionally, we extend our appreciation to the Integrative Genomics of Ageing Group colleagues for their valuable assistance.

## Data availability

To be added.

## Conflicts of interest

JPM is CSO of YouthBio Therapeutics, an advisor/consultant for the Longevity Vision Fund, 199 Biotechnologies, and NOVOS, and the founder of Magellan Science Ltd, a company providing consulting services in longevity science.

GA is CEO of Fagus Antibody services, a company providing custom antibody services to biotech and academia.

